# Switch of TIR signaling by a Ca^2+^ sensor activates ADR1 recognition of pRib-AMP-EDS1-PAD4 for stomatal immunity

**DOI:** 10.1101/2024.10.29.620780

**Authors:** Hanling Wang, Jiaxin Tan, Xiulin Cui, Yuhan Bai, Shang Gao, Brian Staskawicz, Susheng Song, Chuangye Yan, Tiancong Qi

**Affiliations:** Center for Plant Biology, School of Life Sciences, Tsinghua University, Beijing 100084, China; College of Life Sciences, Capital Normal University, Beijing 100048, China; Department of Plant and Microbial Biology, University of California, Berkeley, CA 94720 USA

## Abstract

Plants swiftly close stomata upon detecting pathogen entry, a crucial defense termed stomatal immunity. The process is initiated by cell-surface pattern recognition receptors (PRRs) that perceive pathogen-associated molecular patterns (PAMPs) and evoke a series of early cellular responses including calcium ions (Ca^2+^) influx, and is conducted by the intracellular nucleotide-binding leucine-rich-repeat receptors (NLRs) ADR1s within an EDS1-PAD4-ADR1 module. However, the underlying mechanisms linking PRR signaling to the NLRs ADR1s remain unclear. Here, we show that the *Nicotiana benthamiana* Toll/interleukin-1 receptor (TIR)-only protein Stomatal TIR1 (STIR1) produces the immune molecule pRib-AMP, induces formation of EDS1-PAD4-ADR1 complexes, and mediates stomatal immunity. The Inhibitor of Stomatal Immunity C2-domain protein 1 (ISIC1) interacts with and constrains STIR1 function at basal condition, whereas upon pathogen infection, ISIC1 senses Ca^2+^ signals and de-represses STIR1 signaling. Cryo-electron microscopy structure of pathogen infection-elicited *Arabidopsis* AtEDS1-AtPAD4-AtADR1-L2 complex reveals the pRib-AMP binding to AtEDS1-AtPAD4 receptor and the AtADR1-L2 recognition of pRib-AMP-AtPAD4-AtEDS1 for stomatal immunity. Collectively, this study uncovers a repression/de-repression mechanism linking PRR signaling to NLRs by a Ca^2+^ sensor/TIR-only node, and elucidates an NLR recognition mechanism of the pRib-AMP-EDS1-PAD4 complex in governing innate immunity.

**Synopsis:** At basal condition, the Ca^2+^ sensor ISIC1 interacts with and inhibits the TIR-only protein STIR1; upon pathogen infection, ISIC1 perceives Ca^2+^ signal and releases STIR1 to produce pRib-AMP; the EDS1-PAD4 receptor binds pRib-AMP and is recognized by the NLR ADR1-L2, thereby activating stomatal immunity.

## Introduction

Plants have evolved sophisticated mechanisms to detect pathogen presence, and respond by a series of defense responses including guard cell-specified stomatal closure, to effectively defend pathogen infection (Jones and Dangl, 2006; Melotto et al., 2006). Pathogen molecules are perceived by two types of plant immune receptors. The plasma membrane (PM)-resided cell surface pattern recognition receptors (PRRs) recognize apoplastic pathogen-associated molecular patterns (PAMPs), evoke PAMP-triggered immune responses (PTI) including calcium ions (Ca^2+^) influx as an earliest cellular response (Tian et al., 2019; Tian et al., 2020; Koster et al., 2022; Wang et al., 2024a), and activate stomatal immunity to prevent pathogen entry (Couto and Zipfel, 2016; Liang and Zhou, 2018; Thor et al., 2020; Hou et al., 2024). The intracellular nucleotide-binding leucine-rich-repeat receptors (NLRs) recognize pathogen effectors, and activate effector-triggered immunity (ETI), often accompanied by hypersensitive response (HR) cell death at infection sites restricting pathogen spread (Cui et al., 2015; Jones et al., 2016). PTI and ETI interplay to boost immunity (Ngou et al., 2021; Pruitt et al., 2021; Tian et al., 2021; Yuan et al., 2021).

Plant NLRs harbor a C-terminal leucine-rich-repeat (LRR) domain for perception, a central nucleotide-binding domain shared with APAF-1, various R-proteins and CED- 4 (NB-ARC). According to their N-terminal domains, they are categorized into three major types: Toll/interleukin-1 receptor (TIR)-NLRs (TNLs), coiled-coil (CC)-NLRs (CNLs), and RPW8-type CC-NLRs (RNLs) (Adachi et al., 2019; Feehan et al., 2020; Gong et al., 2023; Maruta et al., 2023). The Arabidopsis CNL AtZAR1 assembles into resistosomes as Ca^2+^ influx channels to mediate HR cell death (Wang et al., 2019a; Bi et al., 2021). By contrast, effector-activated tetrameric TNL resistosomes act as NADase holoenzymes within TIR domains to hydrolyze nicotinamide adenine dinucleotide (NAD^+^) and trigger HR cell death (Horsefield et al., 2019; Wan et al., 2019; Ma et al., 2020; Martin et al., 2020).

RNLs include the Activated Disease Resistance 1 (ADR1) and N Required Gene 1 (NRG1) subfamilies (Grant et al., 2003; Peart et al., 2005; Collier et al., 2011; Jubic et al., 2019; Wang et al., 2024b). *Nicotiana benthamiana* (*Nb*) encodes only one of each (NbADR1 and NbNRG1), whereas *Arabidopsis* has three redundant ADR1 members (AtADR1, AtADR1-like 1 (AtADR1-L1), and AtADR1-L2) and two NRG1s (AtNRG1.1 and AtNRG1.2) (Grant et al., 2003; Peart et al., 2005; Collier et al., 2011; Wang et al., 2024b). The *Arabidopsis* lipase-like protein dimeric receptors AtEDS1- AtPAD4 and AtEDS1-AtSAG101 bind to the ATR1-AtRPP1^TNL^ holoenzyme-generated pRib-AMP/pRib-ADP and ADPr-ATP/di-ADPR, and subsequently form pRib-AMP/- ADP-bound AtEDS1-AtPAD4-AtADR1-L1 complex and ADPr-ATP/di-ADPR-bound AtEDS1-AtSAG101-AtNRG1 complexe, respectively (Huang et al., 2022; Jia et al., 2022). The AtEDS1-AtPAD4-AtADR1 and AtEDS1-AtSAG101-AtNRG1 modules somehow separately activate AtADR1 and AtNRG1 resistosomes at the PM as Ca^2+^ channels to trigger HR cell death (Jacob et al., 2021).

Leaf surface guard cells form stomata and sense pathogens by PM-localized PRRs to initiate stomatal closure (Melotto et al., 2006; Hou et al., 2024; Melotto et al., 2024). Upon perceiving the PAMP flg22 (a peptide derived from flagellin), the PRR complex FLS2-BAK1 and the cytoplasmic kinase BIK1 activate the guard cell Ca^2+^ channel OSCA1.3, and cause Ca^2+^ influx, cytoplasmic Ca^2+^ elevation and stomatal closure (Thor et al., 2020). The intracellular RNL module EDS1-PAD4-ADR1 is essential for PAMP-elicited stomatal immunity (Wang et al., 2024b). However, the molecular mechanisms receiving the PAMP-PRR signaling (e.g., Ca^2+^ signal) and switching on the EDS1- PAD4-ADR1 node remain unknown.

Here, we reveal that a double C2-domain Ca^2+^ sensor and a TIR-only protein create a de-repression mechanism that controls TIR signaling-activated EDS1-PAD4-ADR1- mediated stomatal immunity. Structural analysis of pathogen infection-elicited AtEDS1-AtPAD4-AtADR1-L2 complex emphasizes pRib-AMP as an immune signal for the AtEDS1-AtPAD4 receptor, and elucidates AtADR1-L2 recognition mechanism of pRib-AMP-AtEDS1-AtPAD4 complex. Our findings uncover a receptor signaling cascade and a NLR recognition of pRib-AMP-EDS1-PAD4 complex in stomatal immunity.

## RESULTS

### The TIR-only protein NbSTIR1/2 mediates stomatal immunity via the NbEDS1- NbPAD4-NbADR1 module

TIR domain-only proteins, lacking NB-ARC and LRR domains, are present across plant species (Lapin et al., 2022). However, their inherent roles and mechanisms in defense responses, particularly stomatal immunity, are poorly understood.

In an attempt to explore roles of typical TIR-onlys in stomatal immunity, we discovered that silencing two homologs Nbe03g30820.1 and Nbe18g26860.1 (> 94% amino acid identity), hereafter named Stomatal TIR 1/2 (NbSTIR1/2) (Fig S1), drastically compromised stomatal immunity in *Nb*, as evidenced by susceptibility to spray-inoculation of the bacterial pathogen *Pseudomonas syringae* pv. *tomato* (*Pst*) DC3000 ΔHopQ1 (evade HopQ1-triggered Roq1^TNL^-mediated ETI in *Nb* plants), partial insensitivity to *Pst* DC3000 ΔHopQ1-induced stomatal closure, and increased pathogen entry of the luminescent *P. syringae* pv. *maculicola* (*Psm*) ES4326 into leaf apoplast (Fig. S2). To verify these findings, we generated *Nb stir1 stir2* double knockout gene-editing mutants, and found that they were also compromised in stomatal immunity to *Pst* DC3000 *hrcC^−^* (defective in type III secretion and in triggering ETI) and *Psm* ES4326 (Fig. S3A-B, 1A-C), with a redundant effect of them in *Pst* DC3000 *hrcC^−^*-induced stomatal closure (Fig. S3C). Consistent with their roles in stomatal immunity, *NbSTIR1/2* expression was responsive to *Pst* DC3000 *hrcC^−^* infection (Fig. 1D). These results underscore the crucial roles of *NbSTIR1/2* in mediating stomatal immunity.

**Figure 1.**
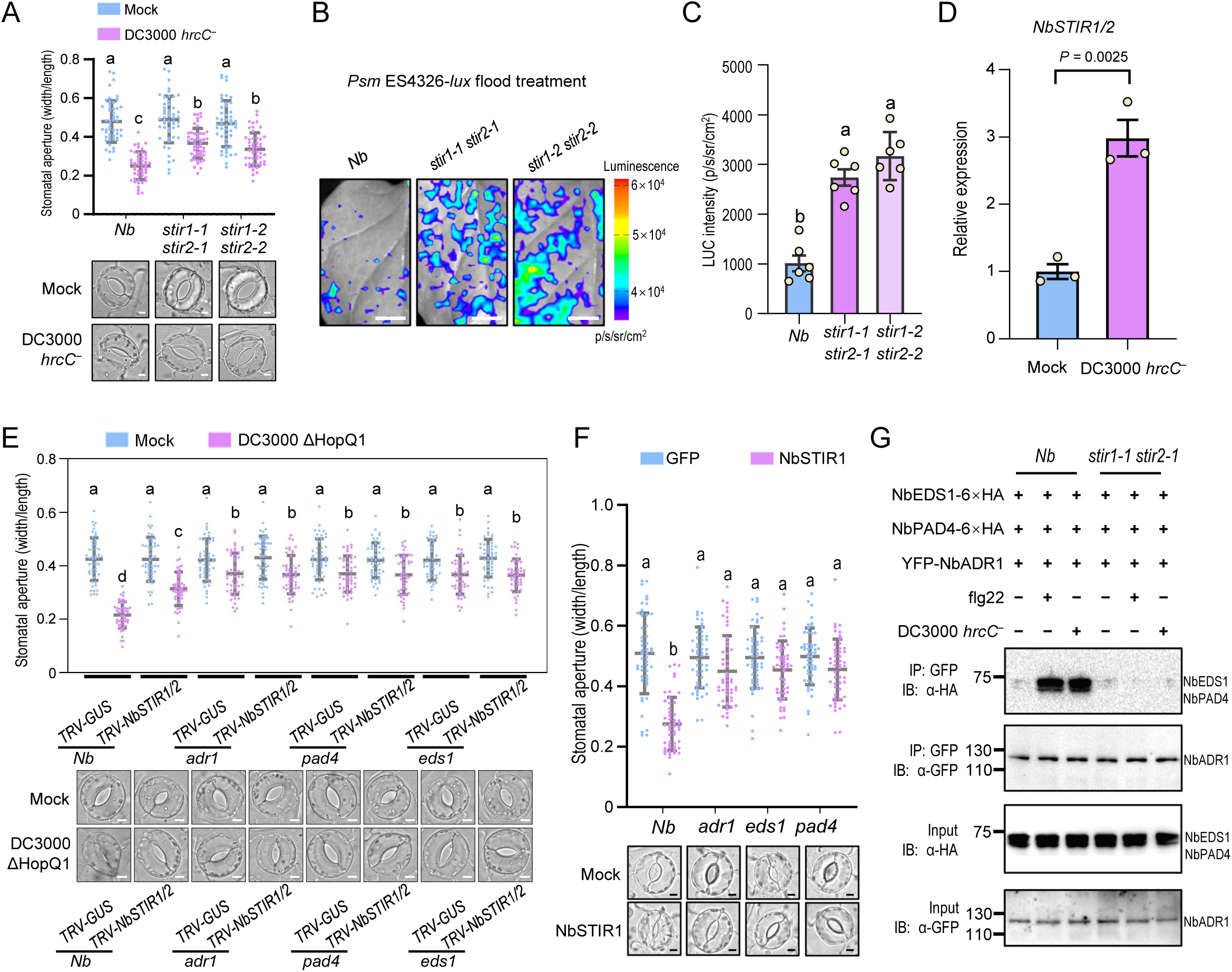
NbSTIR1/2 mediate stomatal immunity via the NbEDS1-NbPAD4- NbADR1 module in *N. benthamiana.* (A) Apertures and images of stomata in wild-type *N. benthamiana (Nb*) and *stir1 stir2* mutants after 1 h of flood treatment without (Mock) or with *Pseudomonas syringae* pv. *tomato* (*Pst*) DC3000 *hrcC*^−^. Data are means ± SD (n = 50). (B-C) Bacterial entry assay showing images (B) and quantification (C) of bacteria in leaves of wild-type *Nb* and *stir1 stir2* mutants. The luciferase (LUC) intensities were recorded by a CCD detector after 1 h of flood treatment with *P. syringae* pv. *maculicola* (*Psm*) ES4326-*lux*. Data are means ± SD (n = 6 independent biological replicates). Scale bars, 1 cm. (D) Quantitative real-time PCR (RT-qPCR) analysis of *NbSTIR1/2* in *Nb* leaves after 2 h of flood treatment without (Mock) or with *Pst* DC3000 *hrcC*^−^. Data are means ± SD (n = 3 independent biological replicates). (E) Stomata apertures in leaves of wild-type *Nb*, *adr1*, *eds1* and *pad4* with *TRV-GUS* or *TRV-NbSTIR1/2* after 1 h of flood treatment without (Mock) or with *Pst* DC3000 ΔHopQ1. Data are means ± SD (n = 50). (F) Stomata apertures in leaves of wild-type *Nb*, *adr1*, *eds1* and *pad4* with transient overexpression of GFP-Flag or NbSTIR1. Data are means ± SD (n = 50). (G) Co-immunoprecipitation (Co-IP) assay showed that *stir1 stir2* compromised *Pst* DC3000 *hrcC*^−^ and flg22-induced association of NbADR1 with NbEDS1 and NbPAD4 in *Nb* leaves. The total proteins were immunoprecipitated with the anti-GFP agarose beads, and the IP products were detected by immunoblotting using anti-GFP or anti-HA antibody. IB, immunoblotting; IP, immunoprecipitation. Data were analyzed by one-way ANOVA by Tukey’s post hoc test (P < 0.05; A, C, E, F) or two-sided Student’s *t*-test (D). Scale bars, 5 μm (A, E, F).

We investigated whether NbSTIR1/2 and the NbEDS1-NbPAD4-NbADR1 module participate in the same pathway of stomatal immunity. Silencing *NbSTIR1/2* did not exacerbate the defect of stomatal closure response in *Nb eds1*, *pad4*, and *adr1* (Fig. 1E). Transient overexpression of NbSTIR1 significantly enhanced stomatal closure in *Nb* wild-type leaves, whereas this effect was impaired in *Nb eds1*, *pad4*, and *adr1* mutants (Fig. 1F). *Pst* DC3000 *hrcC*^−^ infection and flg22 treatment induced formation of the NbEDS1-NbPAD4-NbADR1 complex, while this effect was largely impaired by the *stir1 stir2* mutant (Fig. 1G). These results suggest that NbSTIR1/2 act through the NbEDS1-NbPAD4-NbADR1 module to positively regulate stomatal immunity.

Additionally, overexpression of NbSTIR1/2 triggered HR cell death in *Nb* leaves in a completely or largely *NbEDS1*-dependent manner (Fig. S3D-E). Several other TIR- onlys including AtRBA1 also trigger cell death in an NbEDS1-dependent manner (Nishimura et al., 2017; Wan et al., 2019; Johanndrees et al., 2023).

### NbSTIR1 NADase activity and pRib-AMP production are essential for PAMP-elicited AtEDS1-AtPAD4-AtADR1-L2 assembly and stomatal immunity

TIR domains generally have a conserved catalytic glutamate (E) residue, crucial for NADase activity and triggering HR cell death (Fig. S1B) (Wan et al., 2019; Johanndrees et al., 2023). Modeling by AlphaFold 3 showed that NbSTIR1 also has the conserved residue E109 that positions in the NADase active center of the head-to-tail TIR dimers (Fig. S1B, S4A). NbSTIR1 exhibited *in vitro* and *in vivo* NADase activities (Fig. S3B-G, 2A), with the *Brachypodium distachyon* TIR-only protein BdTIR as a positive control (Wan et al., 2019). Mutating the E109 residue significantly impaired the NADase activity (Fig. S3B-G, 2A), and attenuated NbSTIR1 overexpression-induced stomatal closure and NbSTIR1/2 overexpression-triggered HR cell death (Fig. 2B, S3H). Consistently, application of nicotinamide (NAM), an inhibitor of TIR-NADase activity (Jacob et al., 2023), dramatically attenuated *Pst* DC3000 *hrcC*^−^-triggered stomatal closure (Fig. 2C). These data suggest that the NADase catalytic activity of NbSTIR1 is crucial for stomatal closure.

**Figure 2.**
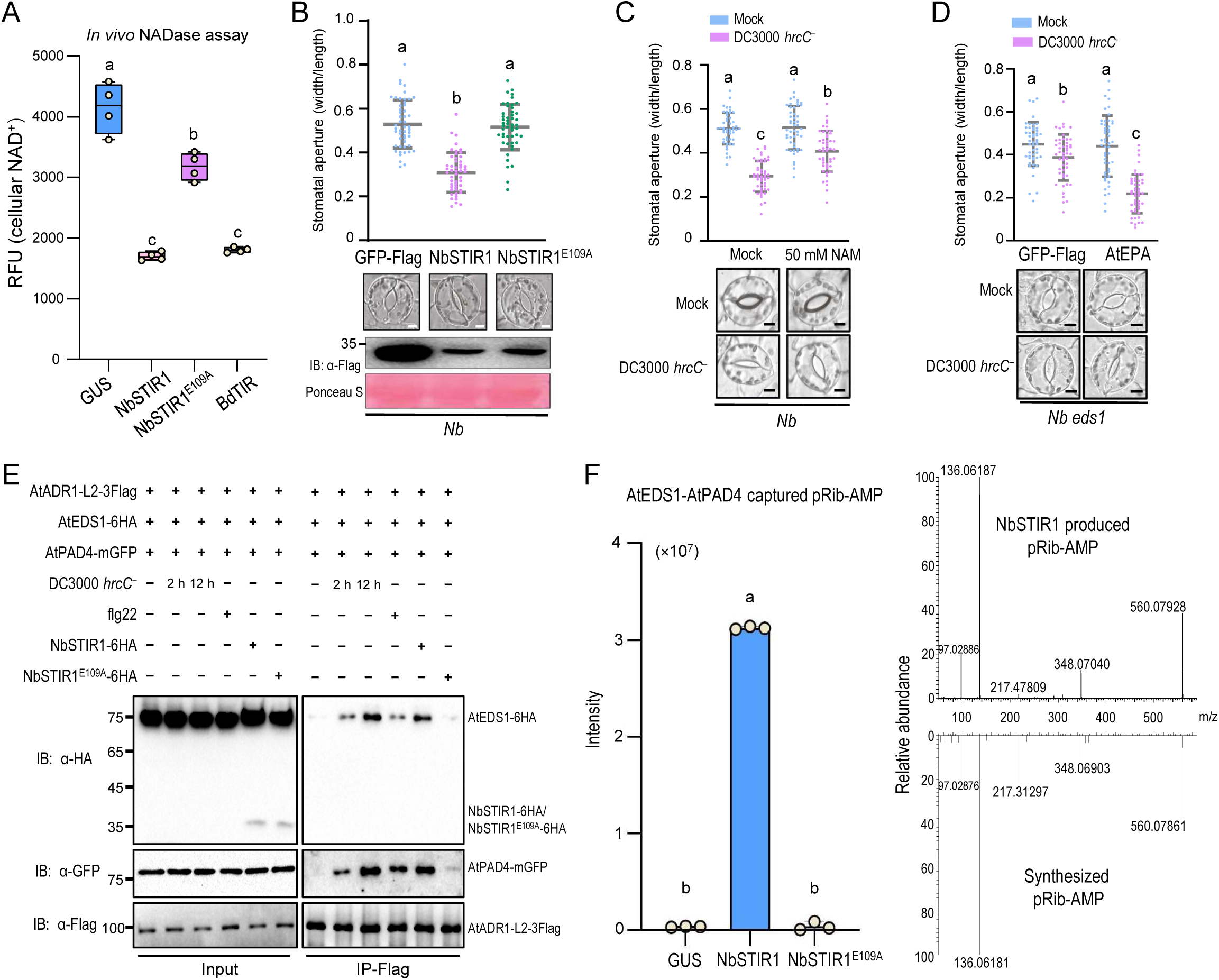
NbSTIR1 NADase activity is required for PAMP-elicited AtEDS1-AtPAD4- AtADR1-L2 assembly and stomatal immunity. (A) Relative NAD^+^ contents in *Nb eds1* leaves at 36h after *Agro-*infiltration of the indicated constructs. RFU, relative fluorescence unit. Data are means ± SD (n = 4 independent biological replicates). (B) Overexpression of NbSTIR1-Flag, but not NbSTIR1^E109A^-Flag, caused stomatal closure in *Nb* leaves. Protein levels in *Nb* leaves were detected by immunoblotting with anti-Flag antibody. Ponceau-S staining of Rubisco was used as a loading control. Data are means ± SD (n = 50). (C) Application of the NADase inhibitor NAM compromised *Pst* DC3000 *hrcC*^−^-induced stomatal closure in *Nb* leaves. Data are means ± SD (n = 50). (D) Co-expression of *Arabidopsis* AtEDS1, AtPAD4 and AtADR1-L2 (AtEPA) restored *Pst* DC3000 *hrcC*^−^-induced stomatal closure in *Nb eds1* leaves. Data are means ± SD (n = 50). (E) Co-IP assay showed that *Pst* DC3000 *hrcC*^−^ infection, flg22, or NbSTIR1 overexpression, but not NbSTIR1^E109A^ overexpression, induced AtADR1-L2 association with AtEDS1/AtPAD4 in *Nb eds1* leaves. The total proteins were immunoprecipitated with the anti-Flag agarose beads, and the IP products were detected by immunoblotting using the anti-HA, anti-GFP or anti-Flag antibody. IB, immunoblotting; IP, immunoprecipitation. (F) Overexpression of NbSTIR1, but not GUS or NbSTIR1^E109A^, generated pRib-AMP that was captured by AtEDS1-AtPAD4 in *Nb eds1* leaves. UPLC-MS/MS analysis of pRib-AMP from AtEDS1-AtPAD4 coexpressed with NbSTIR1, GUS or NbSTIR1^E109A^ (left), and UPLC- MS/MS spectra from AtEDS1-AtPAD4 coexpressed with NbSTIR1 (right top) and synthesized standard pRib-AMP (right bottom) were presented. Data are means ± SD (n = 3 independent biological replicates). Data were analyzed by one-way ANOVA by Tukey’s post hoc test (P < 0.05; A, B, C, D, F). Scale bars, 5 μm (B, C, D).

As the low NbADR1 expression in *Nb* leaves (Fig. 1G) restricted mechanistic dissection, we examined the functional complementation and expression of *Arabidopsis* AtEDS1-AtPAD4-AtADR1-L2 in *Nb* leaves (Fig. 2D-E) to test whether this module could be used for subsequent study. The expression of AtEDS1, AtPAD4, and AtADR1- L2 restored *Pst* DC3000 *hrcC*^−^-induced stomatal closure in *Nb eds1* leaves (Fig. 2D). They formed a complex, with obvious protein levels, in response to *Pst* DC3000 *hrcC*^−^ infection, flg22 treatment, and NbSTIR1 overexpression, but not NbSTIR1^E109A^ overexpression (Fig. 2E). Furthermore, complementary expression of NbSTIR1, but not NbSTIR1^E109A^, restored the *Pst* DC3000 *hrcC*^−^ infection and flg22 treatment-induced interaction of AtADR1-L2 with AtEDS1 and AtPAD4 in the *stir1 stir2* mutant (Fig. S4I). These results suggest that the AtEDS1-AtPAD4-AtADR1-L2 module confers intact functions, including pathogen infection, flg22, and NbSTIR1-induced complex formation, during signaling transduction of stomatal immunity in *Nb* leaves, and that the NADase catalytic activity of NbSTIR1 is essential for AtEDS1-AtPAD4-AtADR1-L2 complex formation and stomatal closure (Fig. 2A-E, S4).

When expressed in insect cells, the ATR1/AtRPP1^TNL^ holoenzyme generates pRib-AMP/pRib-ADP and facilitates AtEDS1-AtPAD4 interaction with AtADR1-L1 (Ma et al., 2020; Huang et al., 2022). By contrast, based on the results of Fig 2A-E, we hypothesize that, during pathogen infection-triggered stomatal closure, the activated NbSTIR1, but not the inactive NbSTIR1^E109A^, produces bioactive molecules that bind to AtEDS1-AtPAD4 and promote their interaction with AtADR1-L2. To test this, we purified AtEDS1-AtPAD4 complex co-expressed with NbSTIR1, NbSTIR1^E109A^, or GUS from *Nb* leaves, respectively, and conducted ultra-performance liquid chromatography-mass spectrometry (MS)/MS (UPLC-MS/MS) to identify NbSTIR1- produced AtEDS1/AtPAD4-captured molecules (Fig. S5A). Results showed that pRib-AMP, but not pRib-ADP, was significantly enriched in the AtEDS1-AtPAD4 sample co-expressed with NbSTIR1, whereas neither pRib-AMP nor pRib-ADP were detected in the other samples (Fig. 2F, S5B-E, S6). These results suggest that pRib-AMP is an NbSTIR1-generated and AtEDS1/AtPAD4-capturable molecule in *Nb* leaves.

Additionally, we observed that *Pst* DC3000 *hrcC*^−^ infection promoted interaction of AtADR1-L2 with AtEDS1 and AtPAD4, but not formation of high molecular weight AtADR1-L2 resistosome triggering HR cell death (Fig. 2E, S7), implying that the AtADR1-L2 resistosome is not involved during stomatal immunity.

In summary, these findings suggest that the NADase activity of NbSTIR1 plays a key role in formation of the AtEDS1-AtAPAD4-AtADR1-L2 complex and stomatal closure, and the NbSTIR1-generated pRib-AMP is an AtEDS1/AtPAD4-capturable molecule in *Nb* leaves.

### The C2-domain protein NbISIC1 interacts with and inhibits NbSTIR1

During exploring roles of pathogen infection-responsive genes in *Nb* stomatal immunity, we found a pair of double C2-domain protein homologs Nbe05g04510.1 and Nbe10g16940.1 with over 98% amino acid identity (Fig. S8). Interestingly, their co-silencing enhanced *Pst* DC3000 ΔHopQ1-induced stomatal closure and resistance to *Pst* DC3000 *hrcC*^−^, and reduced pathogen entry of *Psm* ES4326 (Fig. S9A-E), indicating their inhibitory role in stomatal immunity, and hereafter they were named as Inhibitor of Stomatal Immunity C2-domain protein 1/2 (NbISIC1/2). Double mutants of *NbISIC1/2* were generated to verify this conclusion, and they exhibited increased stomatal immunity to *Pst* DC3000 *hrcC^−^* and *Psm* ES4326 compared with wild-type (Fig. S9F-G, 3A-D), with a redundant role in repressing *Pst* DC3000 *hrcC^—^* infection-induced stomatal closure (Fig. S9H).

We investigated whether NbISIC1/2 suppress stomatal immunity by regulating NbSTIR1 with NbISIC1 as a representative. Co-immunoprecipitation (Co-IP) and pull-down assays showed that NbISIC1 interacted with NbSTIR1 both *in vivo* and *in vitro* (Fig. 3E-F), demonstrating their interaction. A recent study showed that the substrate NAD^+^ induces TIR domains of AtRBA1 and AtRPP1 to undergo phase separation, activate NADase activity, and trigger HR cell death (Song et al., 2024). We observed that NbSTIR1 formed condensates *in vitro* upon NAD^+^ treatment, and NAD^+^-induced NbSTIR1 condensation was gradually repressed by NbISIC1 with a dosage-dependent manner (Fig. 3G-H). Consistently, NbISIC1 compromised the NADase activity of NbSTIR1 (Fig. 3I). *Pst* DC3000 *hrcC*^−^ infection promoted NbSTIR1 condensation in *Nb* leaves, whereas the effect was compromised by NbISIC1 expression (Fig. 3J-K). In support of these findings, NbISIC1 compromised NbSTIR1 overexpression-triggered stomatal closure and HR cell death in *Nb* leaves (Fig. 3L, S9I). These results suggest that NbISIC1 interacts with NbSTIR1, and inhibits the NbSTIR1 condensation, NADase activity, and NbSTIR1 overexpression-triggered stomatal closure.

**Figure 3.**
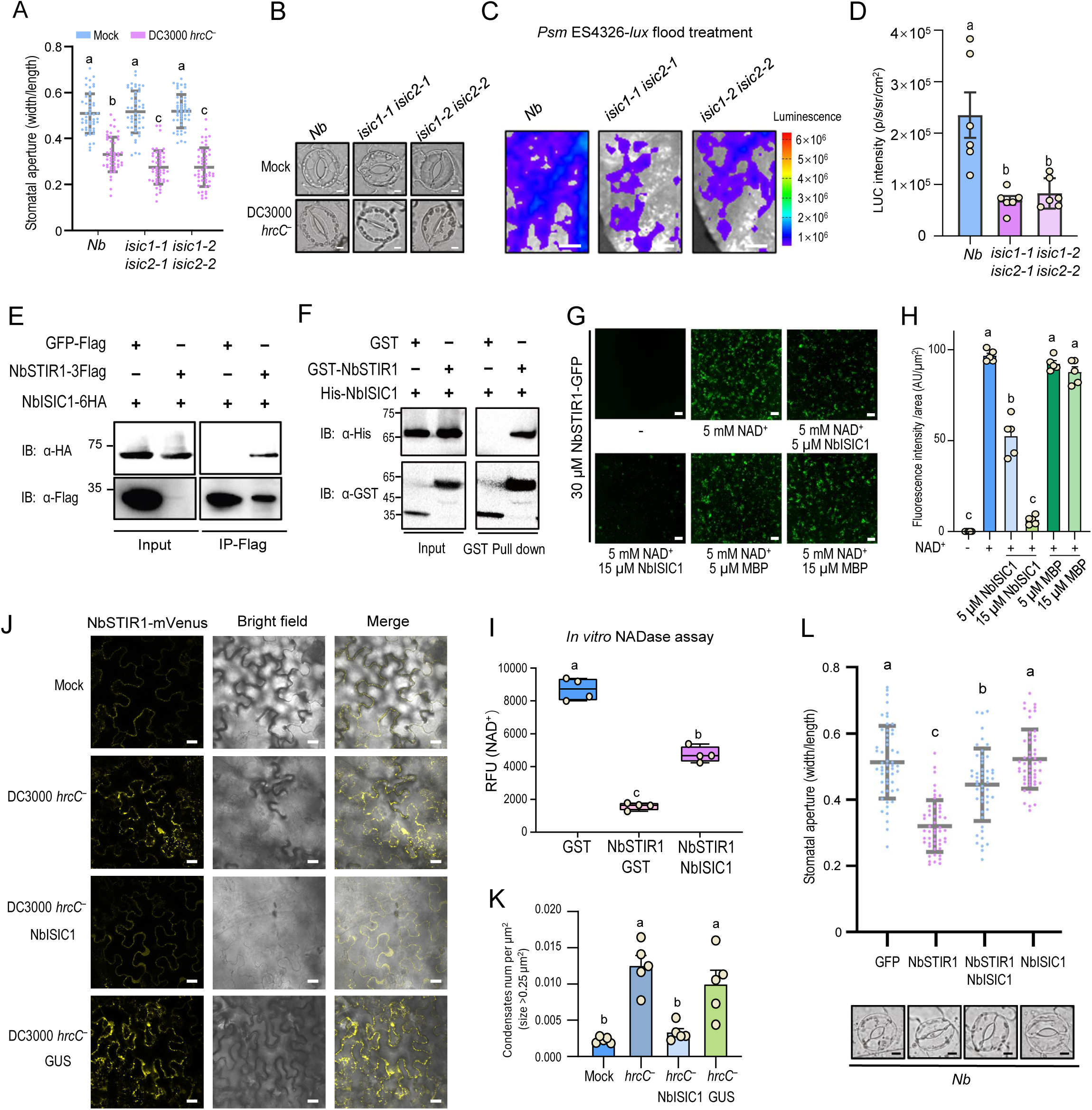
NbISIC1 interacts with and inhibits NbSTIR1 to restrain stomatal immunity. (A-B) Apertures (A) and images (B) of stomata in wild-type *Nb* and *isic1 isic2* mutants after 1 h of flood treatment with mock or *Pst* DC3000 *hrcC*^−^. Data are means ± SD (n = 50). (C-D) Bacterial entry assay showing images (C) and quantification (D) of bacteria in leaves of wild-type *Nb* and *isic1 isic2* mutants. Data are means ± SD (n = 6 independent biological replicates). Scale bars, 0.5 cm. (E) Co-IP assay showed that NbISIC1 associated with NbSTIR1 in *Nb eds1* leaves. (F) Pull-down assay revealed that NbISIC1 interacted with NbSTIR1 *in vitro*. (G-H) Images (G) and quantification (H) of condensates showed that NbISIC1 inhibited the NAD^+^-induced phase separation of NbSTIR1 in a dosage-dependent manner *in vitro*. Data are means ± SD (n = 5). (I) *In vitro* NADase activity assays showed that NbISIC1 inhibits the NADase activity of NbSTIR1. Data are means ± SD (n = 4). (J) Confocal images of *Nb* leaves transiently expressing NbSTIR1-mVenus treated with mock, *Pst* DC3000 *hrcC*^−^, or *Pst* DC3000 *hrcC*^−^ plus expression of NbISIC1 or GUS. Scale bars, 20 μm. (K) Number of NbSTIR1 condensates (size > 0.25 µm²) per µm² in (J). Data are means ± SD (n = 5). (L) Coexpression of NbISIC1 compromised NbSTIR1 overexpression-induced stomatal closure in *Nb* leaves. Data are means ± SD (n = 50). Scale bars, 5 μm. Data were analyzed by one-way ANOVA by Tukey’s post hoc test (P < 0.05; A, D, H, I, K, L); n = independent biological replicates (D, H, I, K).

### NbISIC1 is a Ca^2+^ sensor and relieves NbSTIR1 upon Ca^2+^ signal and pathogen infection

C2-domains typically bind to Ca^2+^ via their aspartic acid (D) residues and mediate Ca^2+^- dependent interactions with phospholipids and proteins (Sutton et al., 1995). As NbISIC1 harbors two C2-domains at the N-terminus (Fig. 4A), we performed surface plasmon resonance (SPR) analysis to examine whether NbISIC1 can bind Ca^2+^. Results showed a high affinity binding of NbISIC1 to Ca^2+^, with a dissociation constant (K_d_) of 33.5 μM (Fig. 4B, S10A). Mutation of the five D residues within C2-domains (NbISIC1^D-penta^) significantly reduced its Ca^2+^ binding ability to a K_d_ of 5.68 mM (Fig. 4C, S10B). These results imply that NbISIC1 is a high-affinity Ca^2+^ sensor.

**Figure 4.**
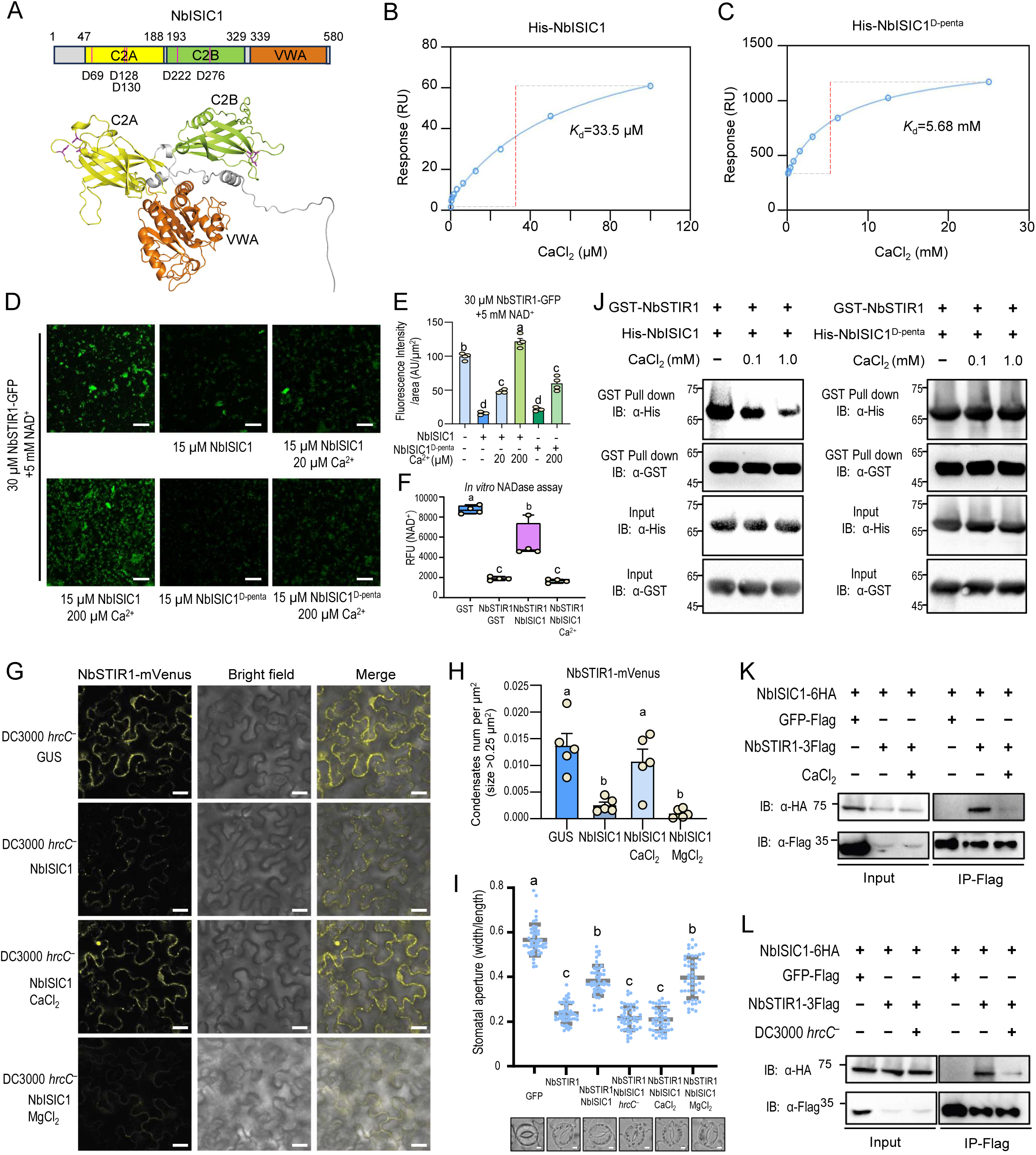
NbISIC1 acts as a Ca^2+^ sensor, and Ca^2+^ signal suppresses NbISIC1 to release NbSTIR1 and trigger its condensation for stomatal immunity. (A) Schematic diagram of NbISIC1 showing its two C2 domains, five aspartic acid (D) residues, von Willebrand A (VWA) domain, and the NbISIC1 structure predicted by AlphaFold3. (B-C) Surface plasmon resonance (SPR) analysis revealed that NbISIC1 bound to Ca^2+^ (B), while the NbISIC1^D-penta^ (D69/128/130/222/276N) mutant compromised binding to Ca^2+^ (C). *K*_d_, dissociation constant. (D-E) Images (D) and quantification (E) of condensates showed that Ca^2+^ application relieved NbISIC1 suppression on NbSTIR1 condensate formation *in vitro*, and the NbISIC1^D-penta^ mutant was resistant to the Ca^2+^ application. Data are means ± SD (n= 4). (F) *In vitro* NADase activity assays showed that Ca^2+^ application relieved the NbISIC1 inhibition on the NADase activity of NbSTIR1. Data are means ± SD (n= 4). (G-H) Images (G) and quantification (H) of NbSTIR1-mVenus condensate showed that Ca^2+^ application de-repressed the NbISIC1 suppression on *Pst* DC3000 *hrcC*^—^induced NbSTIR1-mVenus condensate formation in *Nb* leaves. Data are means ± SD (n= 5). (I) The inhibition of NbISIC1 on NbSTIR1 overexpression-induced stomatal closure was compromised by Ca^2+^ application or *Pst* DC3000 *hrcC*^−^ infection, but not by MgCl_2_, in *Nb* leaves. Data are means ± SD (n = 50). Scale bars, 5 μm. (J) Pull-down assays revealed that the NbISIC1-NbSTIR1 interaction, but not the NbISIC1^D- penta^-NbSTIR1 interaction, was gradually inhibited with increased Ca^2+^ concentration. The protein mixtures were incubated with glutathione agarose resin, and the pull-down products were detected by immunoblotting using anti-GST or anti-HA antibody. (K-L) Ca^2+^ application (L) and *Pst* DC3000 *hrcC*^−^ infection (L) impeded the NbISIC1- NbSTIR1 association in *Nb eds1* leaves. Co-IP assays were performed as in Figure 2E. Data were analyzed by one-way ANOVA by Tukey’s post hoc test (P < 0.05; E, F, H, I); n = independent biological replicates (E, F, H).

Given that Ca^2+^ influx is a prominent feature of earlist cellular responses following PRR activation upon PAMP signals (Tian et al., 2019; Thor et al., 2020), we hypothesize that Ca^2+^ sensing of NbISIC1 de-represses NbSTIR1, thereby facilitating NbSTIR1 activation and stomatal immunity. To test this, we investigated whether Ca^2+^ signal regulates NbISIC1 inhibition on NbSTIR1. We found that Ca^2+^ application dramatically alleviated NbISIC1 suppression on NbSTIR1 condensation, NADase activity, and *Pst* DC3000 *hrcC^−^*-induced NbSTIR1 condensation (Fig. 4D-H). Furthermore, high Ca^2+^ level and *Pst* DC3000 *hrcC*^−^ infection significantly compromised the inhibitory effect of NbISIC1 on NbSTIR1-triggered stomatal closure in *Nb* leaves, compared with Mg^2+^ as a control (Fig. 4I). In contrast, NbISIC1^D-penta^ did not respond to Ca^2+^, and constitutively suppressed NbSTIR1 condensation even under high Ca^2+^ concentration (Fig. 4D-E).

We next investigated whether Ca^2+^ directly impedes the NbISIC1-NbSTIR1 interaction to release NbSTIR1. Pull-down assays showed that Ca^2+^ diminished the NbISIC1-NbSTIR1 interaction in a dosage-dependent manner, but did not impair the NbISIC1^D-penta^-NbSTIR1 interaction (Fig. S10C, 4J). Consistently, Ca^2+^ reduced NbISIC1-NbSTIR1 interaction in *Nb eds1* leaves (Fig. 4K). Accordantly, *Pst* DC3000 *hrcC*^−^ infection attenuated NbISIC1-NbSTIR1 interaction *in planta* (Fig. 4L). These results suggest that upon pathogen infection, elevated Ca^2+^ signals bind NbISIC1, remove NbISIC1 from inhibiting NbSTIR1, and promote NbSTIR1 condensation and activation, facilitating stomatal closure.

### Cryo-EM structure of AtEDS1-AtPAD4-AtADR1-L2 receptor complex

Having shown that the Ca^2+^-NbISIC1-NbSTIR1 signaling mediate stomatal immunity, we next investigated downstream mechanisms of NbSTIR1 particularly the EDS1- PAD4-ADR1 complex formation. Regarding their evident expression and complementary function in stomatal immunity (Fig. 2D-E), we transiently co-expressed AtEDS1, AtPAD4 and AtADR1-L2 in *Nb eds1* leaves with *Pst* DC3000 *hrcC*^−^ infection, and purified the complex using affinity chromatography followed by gel filtration for cryo-EM structural analysis (Fig. S11). Gel filtration analysis showed that co-migration of AtEDS1, AtPAD4, and AtADR1-L2 corresponded to a molecular weight of approximate 220 kDa, indicating the formation of a ternary complex. This complex was subsequently subjected to structural analysis via cryo-EM.

After multiple rounds of 3D classification and refinement using cryoSPARC (Punjani et al., 2017), a map with a global resolution of 2.28 Å was obtained, as determined by gold-standard Fourier shell correlation (Fig. S12, Table S1). The high-quality EM map enabled the assignment of approximately 1,562 side chains within the AtEDS1-AtPAD4-AtADR1-L2 complex, along with their corresponding ligands. The structure showed that AtEDS1, AtPAD4, and AtADR1-L2 assembled in a 1:1:1 ratio (Fig. 5A-B). All domains of AtEDS1 and AtPAD4 are clearly visible. Within the AtADR1-L2 structure, the densities of LRR domain, and helical subdomain 1 (HD1) and winged helix subdomain (WHD) of NB-ARC are evident, while those of CC_RPW8_ and nucleotide-binding subdomain (NBD) of NB-ARC are unvisible (Fig. S13). The LRR of AtADR1-L2 exhibits its characteristic horseshoe shape and anchors to the C-terminal “EDS1-PAD4” (EP domain) of AtPAD4.

**Figure 5.**
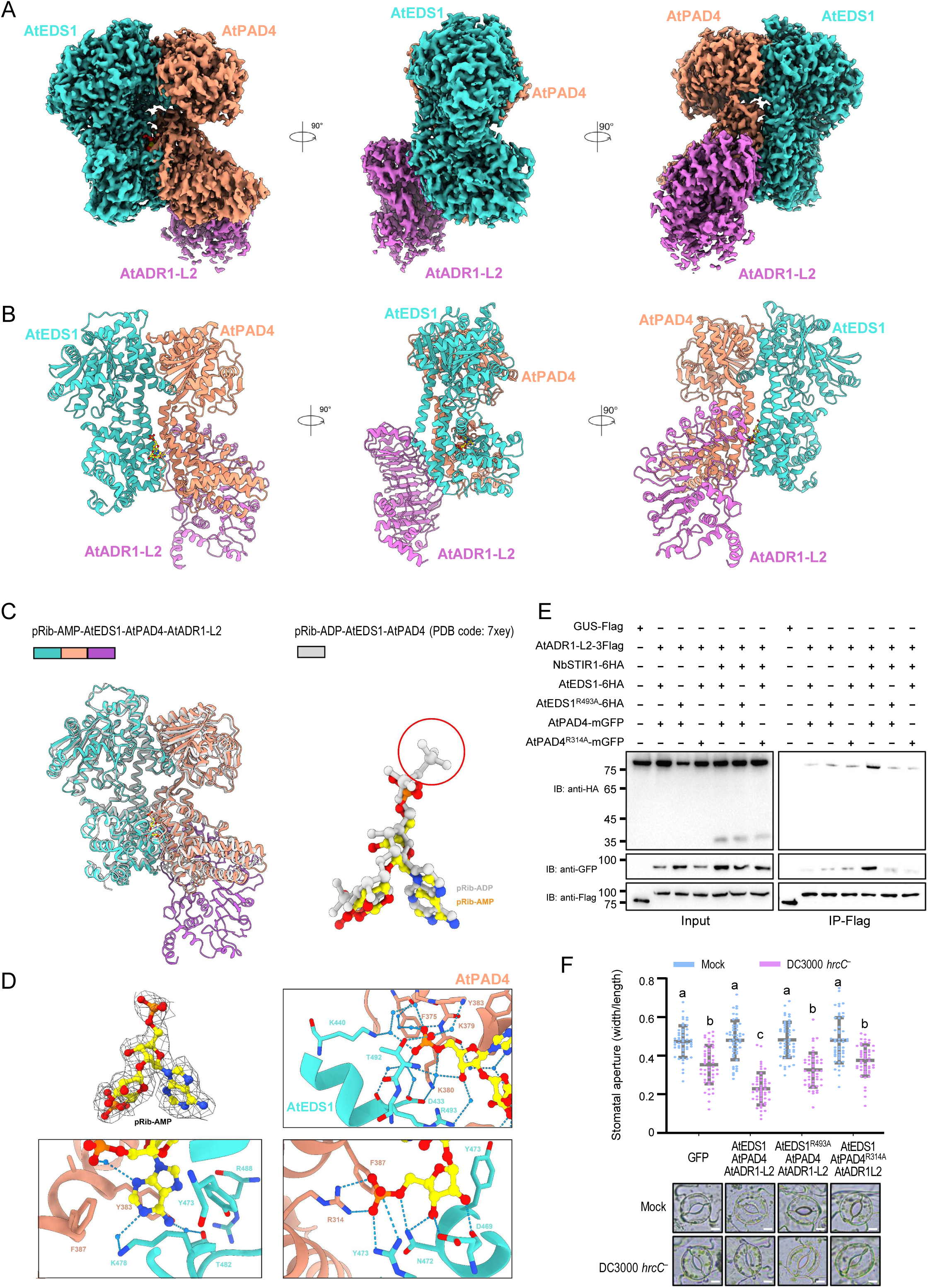
Structure of *Pst* DC3000 *hrcC*^−^ infection-elicited pRib-AMP- AtEDS1/AtPAD4/AtADR1-L2 receptor complex. (A-B) 3D reconstruction (A) and final model (B) of the *Pst* DC3000 *hrcC*^−^ infection-elicited pRib-AMP-bound AtEDS1/AtPAD4/AtADR-L2 ternary complex (PDB: 9JBN) isolated from *Nb eds1* plants in three orientations. The reconstruction and model are shown in the same orientation. (C) Structural alignments between pRib-AMP-bound AtEDS1-AtPAD4-AtADR1-L2 and pRib-ADP-bound AtEDS1-AtPAD4, and between pRib-AMP and pRib-ADP from the determined structures. Structure of pRib-ADP-bound AtEDS1-AtPAD4 is shown in gray. (D) Detailed interactions of pRib-AMP with AtPAD4 and AtEDS1. Blue dashed lines indicate polar interactions. (E) The AtEDS1^R493A^ and AtPAD4^R314A^ mutations within the pRib-AMP binding site disrupted NbSTIR1-induced AtADR1-L2 association with AtEDS1/AtPAD4 in *Nb eds1* leaves. Co-IP assays were performed as in Figure 2E. (F) Co-expression of AtADR1-L2 with the mutant AtEDS1^R493A^ or AtPAD4^R314A^ could not restore *Pst* DC3000 *hrcC*^−^-induced stomatal closure in *Nb eds1* leaves. Data are means ± SD (n = 50). Data were analyzed by one-way ANOVA by Tukey’s post hoc test (P < 0.05). Scale bars, 5 μm.

### pRib-AMP binds to the AtEDS1-AtPAD4 receptor *in planta*

Our cryo-EM structure showed a notable small molecule density within the pocket inbetween the EP domains of AtEDS1 and AtPAD4, and the high resolution of the cryo-EM density map suggests the endogenous molecule as pRib-AMP (Fig. 5A-D). Combined with our UPLC-MS/MS analysis (Fig 2F, S5), these results demonstrate that pRib-AMP is an intrinsic immune molecule for the AtEDS1-AtPAD4 receptor.

In accordance with the pRib-ADP-AtEDS1-AtPAD4 structure (Huang et al., 2022), pRib-AMP also forms an extensive hydrogen bond network including water hydrogen bonds with nearly the same surrounding residues (Fig. 5D), including the pRib moiety with AtEDS1 Asp^469^, Asn^472^ and Tyr^473^, and AtPAD4 Arg^314^, and the AMP moiety with AtEDS1 Lys^478^, Thr^482^, Thr^492^ and Arg^493^. By contrast to the terminal phosphate of pRib-ADP (Huang et al., 2022), water hydrogen bond plays a key role in mediating interaction of the phosphate group of AMP moiety with AtEDS1 Lys^440^, and AtPAD4 Phe^375^ and Lys^379^. These results demonstrate that AtEDS1-AtPAD4 is a receptor complex for pRib-AMP.

To examine the pRib-AMP binding in AtEDS1-AtPAD4-AtADR1-L2 complex, we selectively chose AtEDS1 Arg^493^ and AtPAD4 Arg^314^, critical for pRib-AMP binding, for mutation analysis. Both the AtEDS1^R493A^ and AtPAD4^R314A^ mutations impaired the AtEDS1-AtPAD4-AtADR1-L2 complex formation under NbSTIR1 overexpression, and disrupted *Pst* DC3000 *hrcC*^−^-induced stomatal closure (Fig. 5E-F, S14A), suggesting that the AtEDS1-AtPAD4 binding with pRib-AMP is prerequisite for AtEDS1-AtPAD4-AtADR1-L2 complex formation and *Pst* DC3000 *hrcC*^−^-induced stomatal closure.

### AtADR1-L2 recognition of pRib-AMP-AtEDS1-AtPAD4 receptor complex for stomatal immunity

By comparing structures of apo-AtEDS1-AtPAD4, pRib-ADP-AtEDS1-AtPAD4 (Huang et al., 2022), and pRib-AMP-AtEDS1-AtPAD4-AtADR1-L2 (Fig. 5C, S13B), we found that, upon binding pRib-AMP within a pocket of the AtEDS1-AtPAD4 EP domains, the AtPAD4 EP domain allosterically and counterclockwise rotates ∼10° (Huang et al., 2022) to stretch out and form an interaction interface with AtADR1-L2 LRR domain, and that the overall conformation of AtEDS1-AtPAD4 does not obviously change upon AtADR1-L2 binding, indicating that the AtADR1-L2 binding does not obviously alter the structural integrity of pRib-AMP-AtEDS1-AtPAD4.

The structure showed that AtADR1-L2 acts through its LRR domain and C-terminal loop to bind pRib-AMP-AtEDS1-AtPAD4 by forming two major interfaces (Fig. 5A-B, 6A-B). The AtADR1-L2 LRR domain binds to AtPAD4 EP domain. Within the AtADR1-L2 LRR domain, a series of water mediated hydrogen bonds facilitate interactions with AtPAD4. Two distinct local water clusters, with each composed of three water molecules, form an extensive hydrogen bond network. One cluster bridges AtADR1-L2 His^660^, Asn^684^, His^710^, and AtPAD4 Lys^343^, while the other connects AtADR1-L2 His^660^, Asp^662^, and AtPAD4 Arg^339^ and Tyr^340^. Additionally, AtADR1-L2 Asp^662^ can directly hydrogen bond with AtPAD4 Lys^402^. AtPAD4 Phe^346^ is nestled into a pocket formed by AtADR1-L2 His^710^ and His^660^, further stabilizing the interaction. Consistent with these structural data, the representative mutations AtPAD4^K402A^ and AtADR1-L2^D662A^ in contacting residues of AtPAD4 EP and AtADR1-L2 LRR compromised their capability to induce stomatal closure under *Pst* DC3000 *hrcC*^−^ infection (Fig. 6C, S14B).

**Figure 6.**
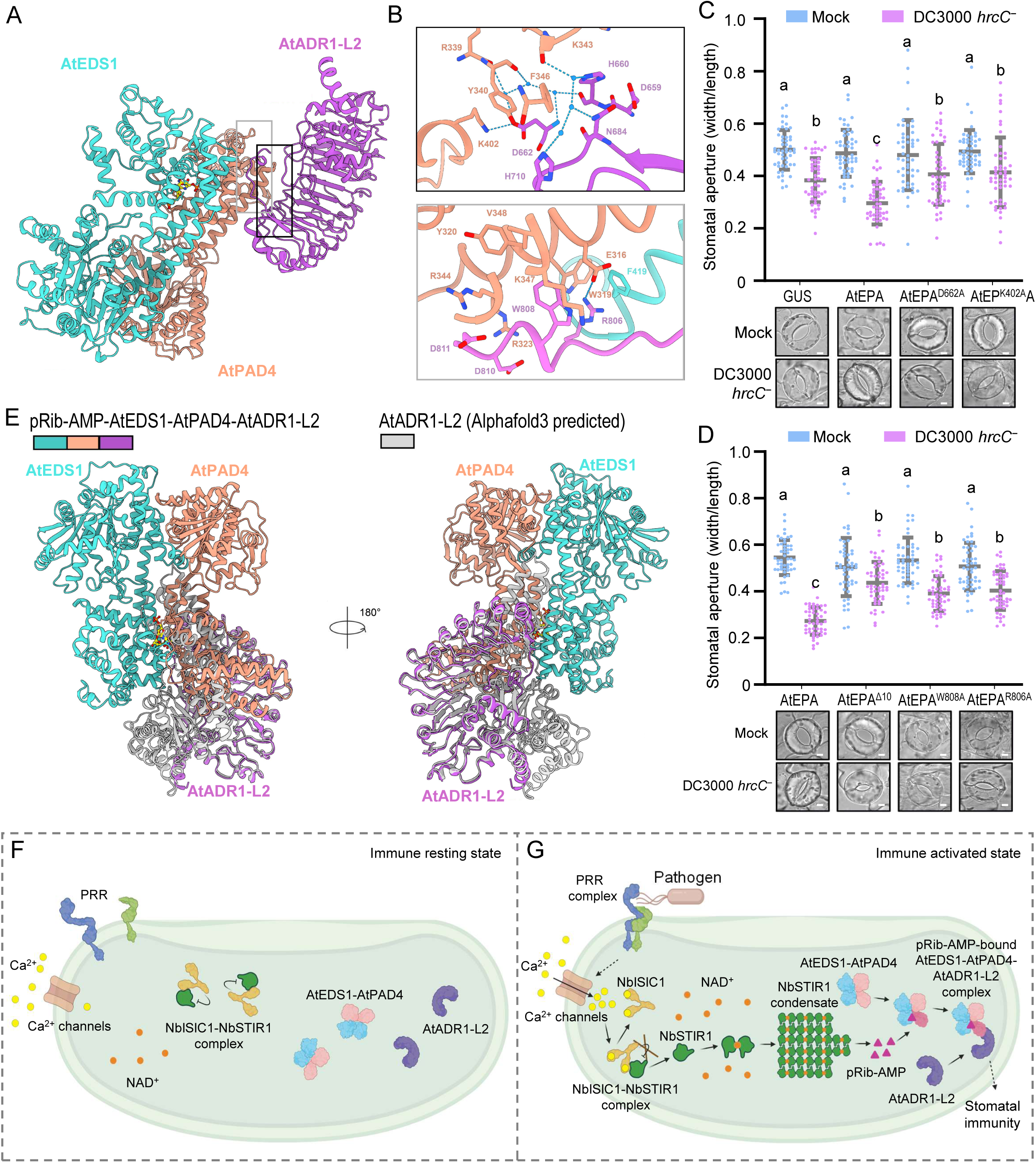
AtADR1-L2 recognition of pRib-AMP-AtEDS1-AtPAD4 for stomatal immunity. (A) Structure showing recognition of pRib-AMP-bound AtEDS1-AtPAD4 by the LRR domain of AtADR1-L2 with two interfaces framed in black and gray respectively. (B) Detailed interactions between AtPAD4 EP domain and AtADR1-L2 LRR domain at their interfaces (black and gray frame in [A]). Blue dashed lines indicate polar interactions. (C-D) Coexpression of AtEDS1/AtPAD4/AtADR1-L2 (AtEPA) with mutations AtPAD4^K402A^ (EP^K402A^A, in C), AtADR1-L2^D662A^ (EPA^D662A^, in C), AtADR1-L2^R806A^ (EPA^R806A^, in D), AtADR1-L2^W808A^ (EPA^W808A^, in D), or deletion of AtADR1-L2 C-terminal 10 amino acids (AtADR1-L2^Δ10^, EPA^Δ10^, in D) was unable to restore *Pst* DC3000 *hrcC*^−^- induced stomatal closure in *Nb eds1* leaves. Data are means ± SD (n = 50). Data were analyzed by one-way ANOVA by Tukey’s post hoc test (P < 0.05). Scale bars, 5 μm. (E) Structural alignment of the pRib-AMP-bound-AtEDS1-AtPAD4-AtADR1-L2 and the Alphafold3-predicted monomeric AtADR1-L2 structure shown in gray. Color codes for AtEDS1, AtPAD4, and AtADR1-L2 are indicated. (F-G) A model of stomatal immunity for this study. At resting condition (F), the Ca^2+^ sensor NbISIC1 interacts with and inhibits the TIR-only protein NbSTIR1, whereas the AtEDS1-AtPAD4 receptor and the RNL receptor AtADR1-L2 remain inactive. Upon pathogen infection (G), guard cell PRRs detect PAMP signals (e.g., flg22), and activate Ca^2+^ influx channels to elevate cytoplasmic Ca^2+^ concentration. NbISIC1 binds to and senses Ca^2+^ signal, and releases NbSTIR1 to bind its substrate NAD^+^, leading to NbSTIR1 condensation, enzyme activation, and production of the immune molecule pRib-AMP. The AtEDS1-AtPAD4 receptor binds the pRib-AMP ligand, causing a conformational change in the AtPAD4 EP domain that allosterically activates its interaction and recognition by the RNL receptor AtADR1-L2, and therefore the activated downstream signaling close stomata to prevent pathogen entry.

The C-terminal loop of AtADR1-L2 inserts in a groove interface formed by both EP domains of pRib-AMP-bound AtEDS1-AtPAD4, establishing extensive interactions (Fig. 6B). Arg^806^ at AtADR1-L2 C-terminus forms a π-π interaction with AtEDS1 Phe^419^, and engages in an electrostatic interaction with AtPAD4 Glu^316^, providing additional stability. The following AtADR1-L2 Trp^808^ engages in hydrophobic interactions with AtPAD4 Lys^347^. AtPAD4 Trp^319^, Tyr^320^, Glu^316^, Val^348^, and Arg^344^ form a hydrophobic pocket, further stabilizing these hydrophobic interactions. The two negatively charged terminal residues Asp^810^ and Asp^811^ of AtADR1-L2 C-terminus form two pairs of electrostatic interactions with AtPAD4 Arg^323^ and Arg^344^, respectively. In support of this structural analysis, the C-terminal 10-AA deletion, and the mutations R806A and W808A within C-terminal loop of AtADR1-L2 impaired *Pst* DC3000 *hrcC*^−^-induced stomatal immune responses (Fig. 6D, S14C). These findings collectively indicate that AtADR1-L2 is the authentic receptor of pRib-AMP-AtEDS1-AtPAD4 complex, and the AtADR1-L2 binding to pRib-AMP-AtEDS1-AtPAD4 by its LRR interface and C-terminal loop is essential for pathogen infection-elicited stomatal immunity.

To gain insights into AtADR1-L2 activation, we compared the monomeric AtADR1-L2 structure predicted by AlphaFold3 (Abramson et al., 2024) with the pRib-AMP-AtEDS1-AtPAD4-AtADR1-L2 structure (Fig. 6E, S13A-B). The monomeric AtADR1-L2 likely exists in a self-inhibited state at basal conditions, with CC_RPW8_ being close to or even contacting LRR domain and occupying the space for LRR recognizing AtPAD4 EP, reminiscent of the inactive NLRs including AtZAR1 (Wang et al., 2019b). The EP domain of allosterically activated AtPAD4 compete for the AtADR1-L2 LRR domain probably with a higher affinity. During pRib-AMP-AtEDS1-AtPAD4 recognition, the C-terminal loop of AtADR1-L2 may insert in the groove of EP domains as a key and undergo some degree of movement, and the LRR domain and HD1/WHD subdomains may cope with minimal structural changes to induce LRR-EP domain binding, whereas CC_RPW8_ and NBD must undertake apparent conformational shift to provide space for the LRR-EP binding and avoid clash with AtEDS1-AtPAD4, thereby derepressing AtADR1-L2 and generating the active pRib-AMP-AtEDS1-AtPAD4- AtADR1-L2 complex for regulating stomatal closure.

## Discussion

Stomatal immunity is a critical early defense of plant innate immune system (Melotto et al., 2006; Melotto et al., 2008; Liu et al., 2022; Hou et al., 2024). Both the cell surface PRR FLS2 and the intracellular RNLs ADR1s mediate stomatal immunity (Melotto et al., 2006; Wang et al., 2024b). flg22 recognition by FLS2 triggers a prominent Ca²⁺ burst as an essential earliest cellular event for stomatal closure (Thor et al., 2020) as well as other PTI responses (Tian et al., 2019; Tian et al., 2020; Koster et al., 2022; Wang et al., 2024a). However, the signal transduction mechanism from PAMP/PRR-triggered cytoplasmic Ca²⁺ elevation, including perception/decoding of Ca^2+^ signals, to the ADR1^RNL^ node remain elusive.

We elucidate a crucial Ca^2+^ signal-primed signal transduction pathway of stomatal immunity (Fig. 6F-G): at basal condition, the Ca^2+^ sensor NbISIC1 interacts with and inhibits the TIR-only protein NbSTIR1; upon pathogen infection or PAMP signals with cytoplasmic Ca^2+^ increase, NbISIC1 perceives Ca^2+^ signals and dissociates from NbSTIR1, and the released NbSTIR1 binds NAD^+^, undergoes condensation and NADase activity to generate the immune molecule pRib-AMP, and therefore the AtEDS1-AtPAD4 receptor binds to pRib-AMP and allosterically attracts AtADR1-L2 recognition, ultimately activating stomatal closure. This natural pathway would provide a long-sought mechanism of perceiving and decoding the PAMP-triggered Ca²⁺ burst during immune responses, not only for stomatal immunity.

TIR-only proteins are present in various plant species (Lapin et al., 2022), and some are transcriptionally responsive to pathogen infection or PAMPs (Tian et al., 2021; Johanndrees et al., 2023). However, their physiological and molecular roles are poorly understood. Here, we demonstrate the pivotal role and regulatory mechanism of the TIR-only NbSTIR1 in stomatal immunity, emphasizing that the pathogen infection-induced NbSTIR1 condensation and NADase activity are essential for stomatal immunity (Fig. 1-4, S1-10). NAD^+^ binding induce TIR condensation/NADase activity for triggering HR cell death (Song et al., 2024). By contrast, we propose a removal-of-repression model as a pivotal mechanism mediating pathogen infection-elicited NbSTIR1 activation: upon pathogen infection with Ca^2+^ increase, the binding of Ca^2+^ to NbISIC1 frees NbSTIR1, allowing NbSTIR1 substrate binding, condensation, and enzymatic activation for stomatal immunity (Fig. 3-4). This reasonable model is reminiscent of the de-repression mechanisms of phytohormone pathways including auxin (Dharmasiri et al., 2005), gibberellin (Ueguchi-Tanaka et al., 2005), and jasmonate (Thines et al., 2007), and strigolactone (Yao et al., 2016). In contrast, this model employs a Ca^2+^ sensor as the repressor of TIR-onlys, and the inhibitory effect is de-repressed by Ca^2+^ signals, whereas the sensory mutant confer dominant repression (Fig. 4-5).

In addition to stomatal closure, we notice that NbSTIR1s overexpression led to HR cell death (Fig. S3D, S4H). Similarly, the TIR-only AtRBA1 triggers HR cell death upon sensing the effector HopBA1 (Nishimura et al., 2017). These findings suggest that TIR-onlys and TIR signaling may play diverse immune roles ranging from the mild stomatal closure to the violent ETI responses, depending on the output intensity of their enzymatic activity and immune status, by input of multilayers of regulation: de-repression by NbISIC1 binding to Ca^2+^ at protein level, substrate-induced condensation, and pathogen infection-responsive expression.

We determined the structure of *Pst* DC3000 *hrcC*^−^-induced pRib-AMP-AtEDS1- AtPAD4-AtADR1-L2 complex isolated from *Nb eds1* plants, and our findings consistently reveal that NbSTIR1-generated pRib-AMP is the stomatal immune signal for the AtEDS1-AtPAD4 receptor, and that pRib-AMP binding to AtEDS1-AtPAD4 receptor is essential for forming AtEDS1-AtPAD4-AtADR1-L2 complex (Fig. 2, 4, S5-6). By contrast, both pRib-ADP and pRib-AMP bind to AtEDS1-AtPAD4 when co-expressed with ATR1-AtRPP1 in insect cells (Huang et al., 2022). This discrepancy may be caused by different cell circumstances. Given that pRib-AMP is produced by both TIR-onlys and TNLs (Fig. 2) (Huang et al., 2022), it may regulate a broad range of responses including stomatal closure and cell death, possibly in a concentration-dependent manner. pRib-AMP is also an efficient inhibitor of prolyl hydroxylase purified from chick embryo (Hussain et al., 1989), indicating its potential diverse functions across kingdoms.

Structural analysis of pRib-AMP-AtEDS1-AtPAD4-AtADR1-L2 complex demonstrates that AtADR1-L2 is the genuine receptor of the pRib-AMP-AtEDS1- AtPAD4 receptor complex (Fig. 5-6, S13). AtADR1-L2 is at a self-inhibited state under basal condition. Allosteric pRib-AMP binding counterclockwise rotates and stretches out AtPAD4 EP domain to interact with AtADR1-L2 LRR domain. During LRR binding to AtPAD4 EP domain of pRib-AMP-AtEDS-AtPAD4 complex, AtADR1-L2 LRR and AtPAD4 EP would undergo an induced-fit interaction after initiating interaction with AtADR1-L2 C-terminal loop, and the cooccurring conformational shift of CC_RPW8_ and NBD make space for the interaction, leading to formation of the active pRib-AMP- AtEDS1-AtPAD4-AtADR1-L2 complex. The structure and function of flexible CC_RPW8_ and NBD in the complex and stomatal immunity are unclear. The transition of CC_RPW8_/NBD from flexibility within pRib-AMP-AtEDS1-AtPAD4-AtADR1-L2 complex to rigidity within AtADR1-L2 resistosome might switch AtADR1-L2 function from the mild stomatal immunity to the fierce HR cell death.

In contrast to the EDS1-SAG101-NRG1 modules that are present in dicots but not in monocots, the EDS1-PAD4-ADR1 modules, TIR-onlys including NbSTIR1 and its homologs such as OsTIR, and NbISIC1 homologs including AtBON1 and OsBON1 are present in both dicots and monocots (Fig. S1, S8) (Collier et al., 2011; Wagner et al., 2013; Lapin et al., 2022; Johanndrees et al., 2023). This ubiquity suggests that they are essential and may confer conservative and broad-spectrum roles in plant innate immunity. The pRib-AMP-AtEDS1-AtPAD4-AtADR1-L2 structure would provide mechanistic reference for ADR1-modules in immune responses of various plant species. The PAMP/PRR - Ca^2+^ signal - Ca^2+^ sensor - TIR-only - pRib-AMP - EDS1/PAD4/ADR1 receptor cascade signaling (Fig. 6F-G) would be a general paradigm for diverse immune responses, not limited to stomatal immunity, in most plant species, and warrants further exploration.

Our study elucidates a Ca^2+^ sensor-TIR only-EDS1/PAD4/ADR1 pathway of stomatal immunity. However, there are still scientific questions remaining to be addressed. In guard cells, whether PAMPs and PRRs somehow parallelly recruit the EDS1-PAD4-ADR1 complex (Pruitt et al., 2021; Tian et al., 2021) to efficiently cooperate with the Ca^2+^-NbISIC1-NbSTIR1 signaling is interesting to be elucidated. The mechanism by which NbADR1 activates downstream signaling including NbWRKY40e to induce stomatal closure remains unclear. Notably, *Pst* DC3000 *hrcC^−^* infection promotes formation of the EDS1-PAD4-ADR1 complex for stomatal immunity, but not the oligomeric ADR1 resistosomes as Ca^2+^ channels triggering cell death. How cells distinguish PTI and ETI signals to control ADR1 output as a ternary complex or resistosome to cope with pathogen infection pressure, and whether and how effectors could manipulate ADR1-mediated pathways warrant further investigation.

## Methods

### Plant materials and growth conditions

The *Nicotiana benthamiana* (*Nb*) wild-type, *eds1*, *adr1*, and *pad4* were described previously (Qi et al., 2018; Wang et al., 2024b). The *stir1 stir2* and *isic1 isic2* mutants were generated in this study by using the CRISPR/Cas9 technology (Fig. S3, S9) (Ge et al., 2017), and the single mutants were segregated. *Nb* plants were grown under a 16 h-light/8 h-dark photoperiod (24℃-26℃).

### Vector construction

Coding sequences of *NbSTIR1/2*, *NbISIC1/2*, *NbEDS1*, *NbPAD4*, *NbADR1*, *AtEDS1*, *AtPAD4*, *AtADR1-L2*, or their derivatives were ligated into the vectors pE1776, pCAMBIA1300, pGEX4T, or pET for fusion with tags including StrepII, Flag or HA, and fragments targeting *NbSTIR1/2* or *NbISIC1/2* for gene silencing were ligated into the TRV2 vector, respectively, through routine molecular procedures. Primers for vector construction are listed in Table S2.

### *Agrobacterium*-mediated transient expression

The *Agrobacterium tumefaciens* (GV3101) strains containing the indicated constructs were cultivated at 28℃ overnight, pelleted by centrifugation, and resuspended in infiltration buffer (10 mM MES, 200 μM acetosyringone, 10 mM MgCl_2_, pH 5.6), kept under dark for 2 h (OD_600_ = 0.1-0.6), and subsequently syringe-infiltrated into *Nb* leaves as *Agro-*infiltration.

### Virus induced gene silencing (VIGS)

The VIGS plants were generated using the TRV-based method (Liu et al., 2002). *Agrobacterium* strains containing *TRV1* and *TRV2* constructs with target fragments (OD_600_ = 0.2) were co-infiltrated into leaves of 2-week-old *Nb* plants to generate VIGS plants.

### Bacterial growth assay

*Pseudomonas syringae* pv. *tomato* (*Pst*) DC3000 ΔHopQ1 cultivated at 28℃ were harvested and resuspended in 10 mM MgCl_2_. The bacterial solution (OD_600_ = 0.4) in 0.02% (v/v) Tween-20 was spray-inoculated onto leaves of 4-week-old *Nb* plants, and at 0 or 6 day later, leaf discs were punched out for bacterial quantification as previously described (Qi et al., 2018).

### Stomatal aperture quantification

Leaves of the *Nb* wild-type, mutants, and VIGS plants were harvested and floated on stomatal opening buffer (50 mM KCl, 10 mM MES-KOH, pH 6.15) with the abaxial surfaces exposed to light for 3 h, and subsequently treated with mock (10 mM MgCl_2_) or a suspension of *Pst* DC3000 *hrcC*^−^ or *Pst* DC3000 ΔHopQ1 (OD_600_ = 0.4) for 1 h. For nicotinamide treatment, 50 mM nicotinamide was added in mock and the bacterial suspension for 1h. For AtEDS1/AtPAD4/AtADR1-L2 or NbSTIR1 overexpression-regulated stomatal aperture, 3-4 leaves from 4-week-old *Nb* plants were syringe-infiltrated and subsequently flooded with *Agrobacterium* strains containing the indicated constructs in a buffer with 0.01% (v/v) Tween-20. At 24 h (for NbSTIR1- related constructs) or 28 h (for AtEDS1/AtPAD4/AtADR1-L2-related constructs) after *Agro*-infiltration, leaves were harvest for stomatal aperture quantification, or for 1 h of treatment with mock, DC3000 *hrcC*^−^ (OD_600_ = 0.4), or 50 mM CaCl_2_ or MgCl_2_ and subsequent stomatal aperture observation. Hypersensitive cell death did not occur in *Nb* leaves at 24 h post *Agro*-infiltration of NbSTIR1. Stomatal apertures (width to length) were captured using a Nikon Ti2-U microscope and measured using ImageJ software.

### Pathogen entry assay

*Nb* leaves were floated on stomatal opening buffer under light for 3 h and subsequently exposed to *P. syringae* pv. *maculicola* (*Psm*) ES4326-luxCDABE (OD_600_ = 0.5; OD_600_ = 0.8 for Fig 3B) for 1 h. Leaves were washed in 0.02% Silwet-77 for 10 s, and photographed by a CCD imaging system (PlantView100) for luciferase (LUC) intensity (p/s/sr/cm2) quantification (Fig. 1C, 3C), or by another CCD imaging system (NightSHADE LB 985) for LUC intensity (cps/cm^2^) quantification (Fig. S1D) of the corresponding leaf areas (∼ 3 cm^2^).

### Hypersensitive cell death assay

*Agrobacterium* suspensions (OD_600_ = 0.5) carrying the indicated constructs were infiltrated into leaves of 4-week-old *Nb* plants. Leaves were harvested at 28 h post-*Agro*-infiltration to detect protein levels by immunoblotting, and imaged under white light and long-wave ultraviolet light at 2 day post-*Agro*-infiltration for hypersensitive cell death symptom observation.

### Real-time quantitative PCR (RT-qPCR) analysis

Total RNA was extracted from *Nb* leaves using RNA-easy^TM^ Isolation Reagent (R701- 01, Vazyme). All-in-One 5×RT MasterMix (G592, abm) was utilized for reverse transcription, and RT-qPCR using BlasTaq 2×qPCR MasterMix (G891, abm) was carried out on a BIO-RAD CFX96^TM^ real-time system with *NbActin* as an internal control (Qi et al., 2018). Primers for RT-qPCR are listed in Table S2.

### Coimmunoprecipitation (Co-IP) assay

*Nb* leaves at 28 h after *Agro*-infiltration of the indicated constructs were syringe- and spray-inoculated with mock, flg22 (1 μM), or *Pst* DC3000 *hrcC*^−^ (OD_600_ = 0.5), and at another 1, 2, or 12 h later, the leaves were collected, homogenized in liquid nitrogen, and suspended in extraction buffer (50 mM HEPES, pH 7.5, 150 mM NaCl, 5 mM MgCl_2_, 0.5% Nonidet P-40, 5% glycerol, 6 mM ß-mercaptoethanol, 10 mM DTT, 1×protease inhibitor cocktail). Samples were centrifuged with 12,000 × g at 4°C for 10 min for three times. The supernatant was transferred and mixed with 8 μL anti-Flag resin (A2220, Sigma) or anti-GFP resin (SA070001, Smart-Lifesciences) and incubated at 4°C for 3 h. The products were washed four times with IP buffer (20 mM HEPES, pH 7.5, 150 mM NaCl, 5 mM MgCl_2_, 0.2% Nonidet P-40, 1% glycerol, 1 mM DTT), resuspended in 3 × Laemmli buffer, boiled for 5 min and centrifuged at 2,000 × g for 5 min. The eluted proteins were subjected to SDS-PAGE and analyzed by immunoblotting.

### Blue native PAGE

Following the blue native polyacrylamide gel electrophoresis (PAGE) protocol (Wittig et al., 2006), the total proteins were extracted from *Nb* leaves with pathogen infection as in the Co-IP assay method section by using modified GTEN buffer (10% glycerol, 100 mM Tris-HCl [pH=7.5], 1 mM Na_2_EDTA, 150 mM NaCl, 5 mM DTT, 0.5% DDM, 1×protease inhibitor cocktail). The extracts were mixed with 5×loading buffer (0.1% Ponceau S, 50% glycerol) and subjected to electrophoresis on a 4 - 13% blue native polyacrylamide gel. The native gel electrophoresis was firstly conducted in cathode buffer (50 mM tricine, 7.5 mM imidazole, 0.02 % CBB G-250) at 150 V for 30 min, and then run in a slightly blue 1/10 cathode buffer (50 mM tricine, 7.5 mM imidazole, 0.002 % CBB G-250) at 150 V for 3 h. After electrophoresis, proteins were transferred to a PVDF membrane (GE Healthcare) using electroblotting buffer (50 mM tricine, 7.5 mM imidazole) at 20 V for 3 h, and subsequently detected by immunoblotting.

### NADase assays

The expression of NbSTIR1, NbSTIR1^E109A^, or BdTIR within pET-His constructs was induced in *E. coli* Rosetta (DE3) cells (Tiangen) by isopropyl β-D-1-thiogalactopyranoside (IPTG) for 4 h at 37℃. Proteins were purified with Ni-NTA agarose resin (ThermoFisher), concentrated to approximately 50 μM in 100 mM NH_4_HCO_3_ (MREDA) buffer (pH 7.6). Each NADase reaction mixture (100 μL) contained 100 mM NH_4_HCO_3_ (pH 7.6), 100 μM NAD^+^ (NEB), 10 mM MgCl_2_ (Sigma), 1 mM 3-isobutyl-1-methylxanthine (IBMX, Sigma), 1 mM DTT (Sigma), and either 25 μM TIR proteins or buffer as a control, and were incubated at 25℃ for 16 h. Samples were centrifuged at 12,000 rpm for 10 min, freeze-dried, dissolved in 50 μL of 80% methanol, and analyzed by liquid chromatography.

The NAD^+^ degradation assays were also conducted in a reaction buffer containing 92.4 mM NaCl and 0.64× phosphate-buffered saline. A 50 μL reaction mixture was prepared with final concentrations of 30 μM NbSTIR1, 30 μM NbISIC1 and/or GST, and 5 μM NAD^+^ (NEB) without or with 1mM CaCl_2_. The mixture was incubated at 25°C for 2 h. After incubation, the sample was used to measure the remaining NAD^+^ using the Amplite NAD^+^ Assay Kit (15280, AAT Biosciences).

Three 8-mm discs of *Nb eds1* leaves at 36 h after *Agro-*infiltration of the indicated constructs were harvested into 1.5-mL tubes with a single glass bead and were flash-frozen in liquid nitrogen. The tissues were homogenized using a QIAGEN TissueLyzer II at 30 Hz for 30 seconds. The homogenates were resuspended in 400 mL of ice-cold lysis buffer with 50 mM Tris-HCl (pH 7.5), 150 mM NaCl, 5 mM EDTA, 0.2% Triton X-100, and 10% (v/v) glycerol, and were stored on ice. The lysates were centrifuged at 4°C for 10 min at 5000 rpm. The supernatant was diluted 1:3 in lysis buffer before being added to the Amplite NAD^+^ Assay Kit (15280, AAT Biosciences). The colorimetric reagent was developed for 20 min at room temperature. Colorimetric detection of NAD^+^ was performed in 96-well black-bottom plates (Costar) using a Spark Multimode Microplate Reader (Tecan) with excitation at 420 nm and emission at 480 nm, and the results were expressed as relative fluorescence units.

### *In vivo* STIR1-catalyzed small molecule capture assay

AtEDS1-3×Flag and AtPAD4-3×Flag were co-expressed with GUS, NbSTIR1-6HA or NbSTIR1^E109A^-6HA in *Nb eds1* leaves through *Agro*-infiltration. 10 g leaves were harvested at 28 h post-*Agro*-infiltration. The AtEDS-AtPAD4 complex was purified using anti-Flag resin (A2220, Sigma). The IP products were eluted in IP buffer containing 300 mg/L 3×Flag peptide and concentrated to 20 μL in 100 mM NH_4_HCO_3_ (MREDA) buffer (pH 7.6). Each sample was denatured with 80 μL methanol, incubated at −80℃ for 30 min, and analyzed by liquid chromatography.

### Pull-down assay

The *E. coli* Rosetta (DE3) cells containing the pGEX-4T-GST or pET-His construct with NbSTIR1, NbISIC1, their mutants, or the GST control were cultured to OD_600_ = 0.8 at 37℃. GST and GST-NbSTIR1 were induced with 0.4 mM IPTG for 4 h at 37℃ and purified using Glutathione agarose resin (ThermoFisher). NbISIC1 and its mutant were induced with 0.1 mM IPTG for 16 h at 12℃ and purified using Ni-NTA agarose resin (ThermoFisher). Proteins were concentrated to 50 μM in pull-down buffer containing 20 mM HEPES (pH 7.5), 150 mM NaCl, 0.02% NP-40, 1 mM DTT as input. GST and GST-NbSTIR1 were initially added to 10 μL Glutathione agarose resin (ThermoFisher) and incubated for 2 h at 4℃, and NbISIC1 and its mutant were added to the resin and incubated for additional 2 h or overnight at 4℃, respectively. Each resin was washed for 4 times with the pull-down buffer. For Ca^2+^ gradient treatment, the pull-down buffer containing different concentrations of CaCl_2_ (0, 0.1, 0.5, and 1 mM) were added to equally divide NbSTIR1-NbISIC1/NbISIC1^D-penta^ resin and incubated for 1 h at 4℃. Input and resin were subsequentially boiled in 3×Laemmli buffer for 5 min, centrifuged and analyzed by immunoblotting.

### Surface plasmon resonance (SPR) assay

The kinetics of Ca^2+^ binding to NbISIC1 and its mutant, purified by Ni-NTA agarose resin, were analyzed using a Biacore 8K plus (Cytiva) in filtered and degassed SPR buffer containing 50 mM HEPES (pH 7.5) and 150 mM NaCl at 25℃. NbISIC1 and its mutant were respectively immobilized on CM5 sensor chips, achieving an approximate response unit (RU) of 1000. CaCl_2_ (Sigma) was serially diluted (0.09765625–50 μM for NbISIC1, and 0.09765625–50 mM for NbISIC1^D-penta^) and flowed through the immobilized NbISIC1 and its mutant. The binding signals were recorded by a multi-cycle method with 90 s of association and 90 s of dissociation at a 30 μL/min flow rate. The binding data were analyzed and fitted to a one-site specific binding mode curve using Graphpad Prism. The Ca^2+^ binding constants (*K*_d_) for NbISIC1 and its mutant were calculated from the fitting curve.

### *In vivo* and *in vitro* phase condensation assays

The suspension of *Agrobacterium* strains (OD_600_ = 0.5) harboring *pNbSTIR1-NbSTIR1- mVenus* alone or together with NbISIC1 or GUS were co-infiltrated into *Nb eds1* leaves, and at 28 h later, the leaves were sprayed with mock or *Pst* DC3000 *hrc*C^−^ (OD_600_ = 0.4), and at another 2 h later, the leaves were observed under a microscope. For Ca^2+^ or Mg^2+^ treatment, leaves were soaked in 50 mM CaCl_2_ or MgCl_2_ for 30 min, and then were observed under a microscope. The MBP-6His-NbSTIR1-GFP proteins, purified by Ni-NTA agarose resin, were cleaved by His-TEV protease at 4 °C overnight. The reaction mixture, containing 30 µM NbSTIR1-GFP, 5 mM NAD^+^, and 5 or 15 µM NbISIC1, 15 µM NbISIC1^D-penta^, or 5 or 15 µM MBP, was prepared in a reaction buffer with 25 mM Tris-HCl (pH 8.0), 50 mM NaCl and 5% PEG4000. For Ca^2+^ treatment, NbSTIR1-GFP and NbISIC1 or NbISIC1^D-penta^ were mixed and added to the reaction buffer with 20 or 200 µM CaCl_2_, and the mixtures were incubated at 25 ℃ for 1 h. After incubation, the samples were loaded onto 384-well glass bottom plates (Cellvis) and observed under a confocal microscope.

### Confocal imaging and fluorescence analysis

GFP and mVenus proteins were excited with peaks of 488 nm and 514 nm, respectively, and their emissions between 490 and 530 nm, and between 530 and 560 nm were captured, respectively, by a Zeiss laser scanning microscope (LSM 880). To determine the numbers of condensed NbSTIR1, ImageJ software and Otsu’s method were used to remove background signals and present GFP signals, and figure out results of areas and the corresponding intensities. The “analyze particles” function was used with parameters of size (0.25 µm^2^–infinity) and circularity (0.2–1) to perform particle analysis.

### Ultra-performance liquid chromatography (UPLC) -MS/MS analysis

The UPLC system was coupled with a Q-Exactive HFX orbitrap mass spectrometer (Thermo Fisher, CA) equipped with a heated electrospray ionization (HESI) probe. 2 μL sample was loaded onto a BEH Amide 100 × 2.1 mm column (Waters) for positive mode analysis. Samples were eluted with aqueous solvent containing 5 mM ammonium acetate as eluent from 50% to 98% within 8 min for NAD^+^ or 5 min for pRib-AMP. The stationary phase consisted of 95% acetonitrile with 5 mM ammonium acetate. Data with mass ranges of m/z 300-1500 (NAD^+^) or 300-1200 (pRib-AMP and pRib-ADP) at positive ion mode with data dependent MS/MS acquisition. The full scan spectra and the fragment spectra were collected with resolution of 60,000 and 15,0000, respectively. Source parameters were set as follows: spray voltage, 3000 V; capillary temperature, 320 ℃; heater temperature, 300 ℃; sheath gas flow rate, 35; auxiliary gas flow rate, 10. Data analysis was performed using Xcalibur 4.4 software (CA, Thermo Fisher), and data visualization was conducted using Graphpad Prism.

### Reconstitution and purification of the AtEDS1-AtPAD4-AtADR1-L2 complex

At 22 h after *Agro-*infiltration of StrepII-AtEDS1 (OD_600_ = 0.1), StrepII-AtPAD4 (OD_600_ = 0.1) and Flag-AtADR1-L2 (OD_600_ = 0.6) into *Nb eds1* leaves, *Pst* DC3000 *hrcC*^−^ was spray-inoculated onto the leaves, and another 6 h later, 100 g leaves were collected at 28 h post-inoculation, homogenized and resuspended in 160 mL extraction buffer containing 50 mM HEPES (pH 7.5), 150 mM NaCl, 5 mM MgCl_2_, 0.5% DDM, 5% glycerol, 10 mM DTT, 6 mM β-mercaptoethanol, and 1×protease inhibitor cocktail. The homogenate was centrifuged at 20,000 × g for 1 h, and the supernatant was filtered with miracle cloth and then centrifuged again at 20,000 × g for 1 h. 600 μL anti-Flag resin (Sigma) was added into the supernatant, and the mixture was incubated at 4℃ for 3 h. The resin was washed 5 times of 5 min with wash buffer containing 20 mM HEPES, pH 7.5, 150 mM NaCl, 5 mM MgCl_2_, 0.02% DDM, 1% glycerol, 1 mM DTT. The proteins were eluted twice by incubating with wash buffer containing 300 μg/mL 3×Flag peptide for 12 h at the first time and 2 h at the second time. The eluates were concentrated to 100 μL, and subjected to gel filtration using a Superdex 200 5/150 column. Gel filtration fractions were analyzed by SDS-PAGE and silver-staining. Ideal fractions corresponding to approximately 200 kD, containing AtEDS1, AtPAD4 and AtADR1-L2, were pooled and concentrated to 0.9 mg/mL for cryo-EM sample preparation. For negative staining, the protein was diluted to 0.018 mg/mL.

### Negative-stain electron microscopy

6 μL of purified AtEDS1-AtPAD4-AtADR1-L2 complex was applied to freshly glow-discharged copper grids and incubated for 1 min. 6 μL of 2% (w/v) uranyl acetate dihydrate was utilized to wash the grid for 1 min for twice. The grid was blotted and captured by the FEI Tecnai Spirit transmission electron microscope (Thermo Fisher Scientific).

### Cryo-EM sample preparation and data acquisition

4 μL AtEDS1-AtPAD4-AtADR1 complex purified from *Nb* leaves was applied to glow-discharged Quantifoil holey carbon grids (Quantifoil Cu R1.2/1.3+2 nm C, 300 mesh). The protein was concentrated to 0.9 mg/mL prior to freezing. After application, the grids were blotted for 3.0 seconds under 100% humidity at 4°C and plunge-frozen in liquid ethane cooled by liquid nitrogen using a Vitrobot (Mark IV, Thermo Fisher Scientific). Cryo-EM data were collected on a 300 kV Titan Krios G4 microscope equipped with a Falcon 4 electron detector and a GIF Quantum energy filter (slit width 10 eV). The defocus values ranged from −1.4 to −1.8 μm. Each stack of 32 frames was exposed for a total of 2.56 seconds, with each frame having an exposure time of 0.08 seconds. Micrographs were automatically acquired using EPU software in super-resolution counting mode with a binned pixel size of 0.96 Å. The total electron dose for each stack was approximately 50.56 e-/Å². All 32 frames within each stack were aligned and summed using the whole-image motion correction software, MotionCor2 (Zheng et al., 2017).

### Image processing and 3D reconstruction

All dose-weighted micrographs were manually inspected and imported into cryoSPARC (Punjani et al., 2017). Micrographs with an estimated CTF resolution worse than 4 Å were excluded during exposure curation. CTF parameters were estimated using the patch-CTF method. The Blob picker was initially used to generate good templates via 2D classification, after which the Template picker was employed for particle picking. Particles were extracted with an original box size of 300 pixels and cropped to 128 pixels to speed up calculations. Initial good and bad references were generated using particles from the dataset. For this dataset, 7,126,787 particles were extracted from 14,113 micrographs. After several rounds of 3D classification, 1,779,204 particles were selected for subsequent non-uniform refinement and local refinement, resulting in a final resolution of 2.28 Å.

### Cryo-EM model building and refinement

The initial model of the AtEDS1-AtPAD4-AtADR1-L2 complex was generated using AlphaFold 3 (Abramson et al., 2024). The predicted model was fitted into the cryo-EM map using UCSF ChimeraX (Pettersen et al., 2021). COOT was used for manual adjustment and rebuilding of the structure (Emsley and Cowtan, 2004). Ligand restraint files for refinement were generated using phenix.elbow (Afonine et al., 2018; Liebschner et al., 2019). The final AtEDS1-AtPAD4-AtADR1-L2 model was refined against the corresponding maps using PHENIX in real space with secondary structure and geometry restraints (Afonine et al., 2018). Model overfitting was monitored by refining the model against one of the two independent maps from the gold-standard refinement process and testing the refined model against the other map (Amunts et al., 2014). The structure was validated by assessing Clash scores, MolProbity scores, and Ramachandran plot statistics using PHENIX. Cryo-EM data collection, refinement and validation statistics are summarized in Table S1.

### Accession numbers

The accession numbers for genes mentioned in this study are as follows: *NbSTIR1* (Nbe03g30820.1/Niben101Scf00180g07008.1), *NbSTIR2* (Nbe18g26860.1/Niben101 Scf02391g00012.1), *NbISIC1* (Nbe05g04510.1/Niben101Scf06621g02031.1), *NbISIC2* (Nbe10g16940.1/Niben101Scf03548g02007.1), *NbADR1* (Niben101 Scf02422g02015.1), *NbEDS1* (Niben101Scf06720g01024.1), *NbPAD4* (Niben101Scf02544g01012.1), *AtADR1* (AT1G33560), *AtADR1-L2* (AT5G04720), *AtEDS1* (AT3G48090/*AtEDS1a*), *AtEDS1b* (AT3G48080), and *AtPAD4* (AT3G52430).

### Quantification and statistical analysis

Data are presented as mean ± SD, and the statistical analyses were performed by one-way analysis of variance (ANOVA) followed by Tukey’s post hoc test or two-sided student *t*-test.

## Data availability

All data are provided in the main figures and supplemental figures. Atomic coordinate of AtEDS1-AtPAD4-AtADR1-L2 complex (9JBN) was deposited in the Protein Data Bank (http://www.rcsb.org). The cryo-electron microscopy map (EMD-61320) was deposited in the Electron Microscopy Data Bank (https://www.ebi.ac.uk/pdbe/emdb/). All other data that support the findings of this study and materials used in this study are available from the lead contact upon reasonable request, Tiancong Qi.

## Acknowledgement

We thank Prof. Jijie Chai, Dr. Shijia Huang for providing materials and helpful suggestions, thank Lina Xu, Zi Yang, Kaile Huang, and Yuxuan Zou for their technical assistance or materials. We thank the Ministry of Science and Technology-Key Research and Development Program (2021YFA1301800), the National Natural Science Foundation of China (32370756), Tsinghua University Initiative Scientific Research Program, and the Tsinghua-Peking Center for Life Sciences for funding support.

## Author contribution

T.Q., C.Y., and S.S. conceived and designed the project; H.W., J.T., X.C., and Y.B. performed the experiments. T.Q., C.Y., H.W., J.T., X.C., and S.G., analyzed data and prepared figures and tables. S.S., and B.S. analyzed data, T.Q. and S.S. wrote the manuscript with assistance from all authors. All authors provided comments and contributed to the manuscript preparation.

## Competing interests

The authors declare no competing interests.

## Supplemental Figures and Tables

**Figure S1.**
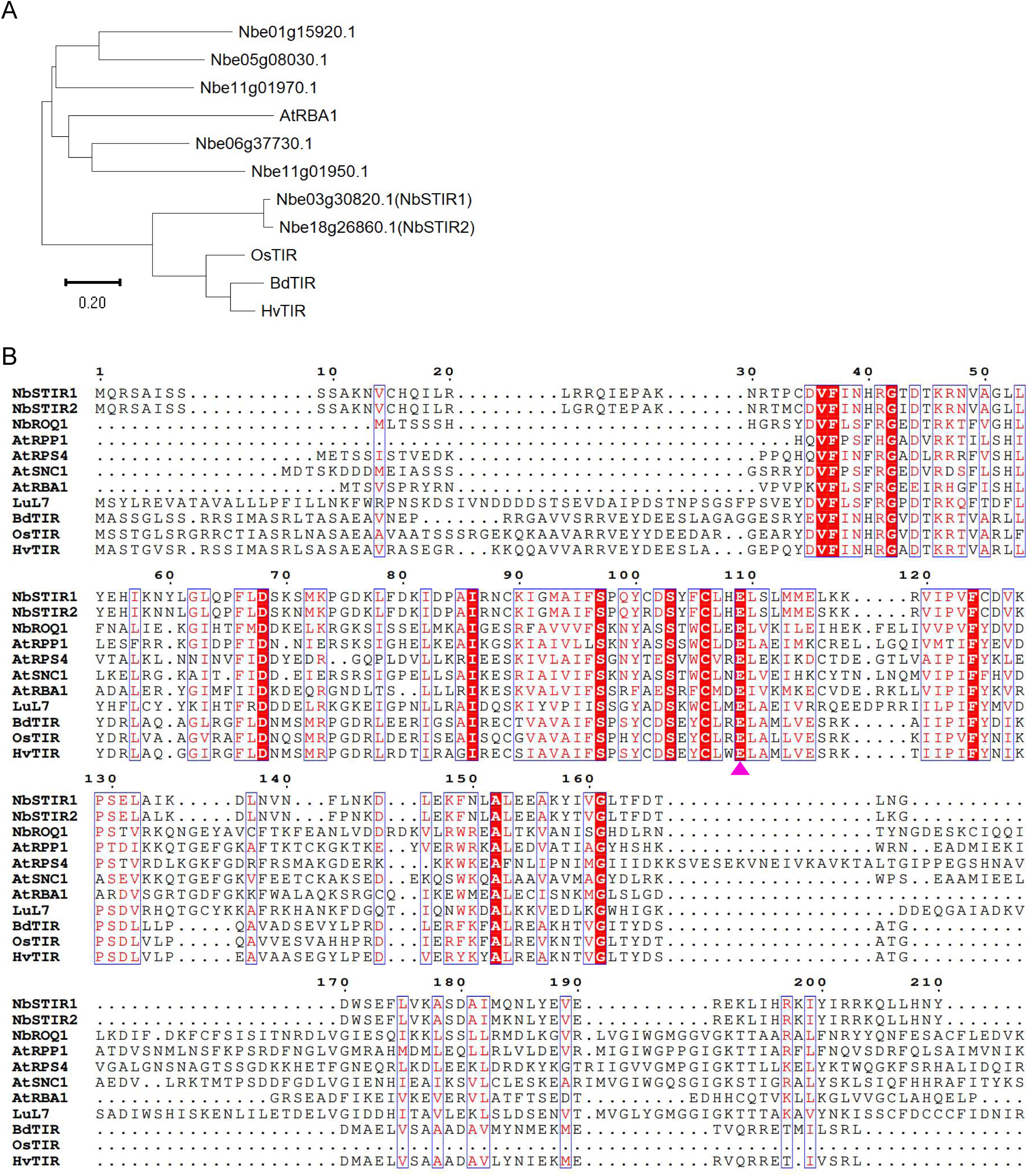
Phylogenetic analysis and protein alignment of NbSTIR1/2 and other TIR-only proteins. (A) Phylogenetic analysis of the TIR-only proteins NbSTIR1, NbSTIR2, other identified TIR-only proteins from *N. benthamiana* (*Nb*), *Oryza sativa* (OsTIR, Os07G0566800), *Hordeum vulgare* (HvTIR, HORVU2Hr1G039670), *Brachypodium distachyon* (BdTIR, XP_003560074.3), and *Arabidopsis* (AtRBA1, AT1G47370). (B) Multiple protein sequence alignment of the TIR-only proteins NbSTIR1, NbSTIR2, OsTIR, HvTIR, BdTIR, AtRBA1, and TIR domains of TNLs from *Nb* NbRoq1, Arabidopsis AtRPP1, AtANC1, and *Linum usitatissimum* LuL7. The conserved E residues for the NADase activity are marked by triangles.

**Figure S2.**
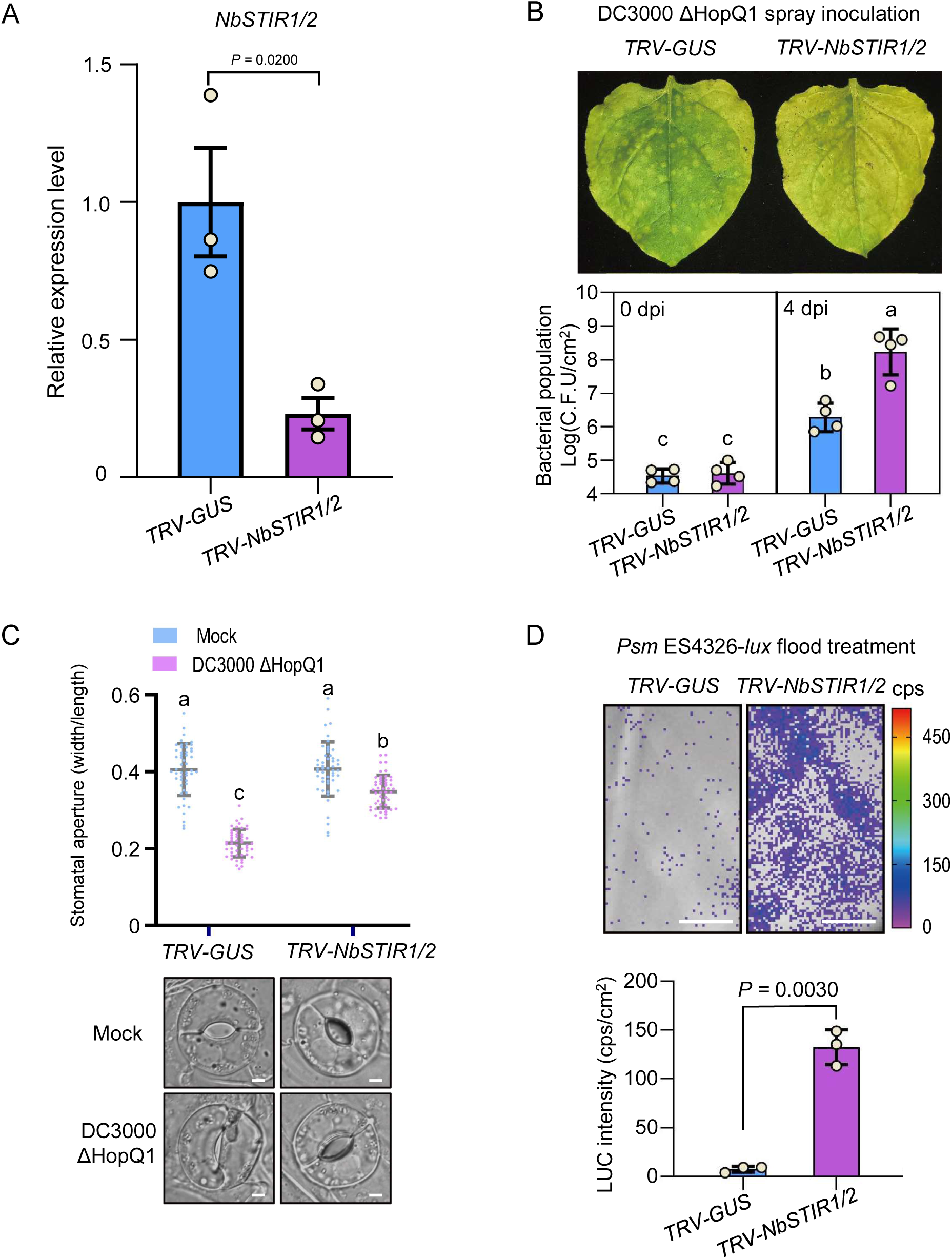
Silencing *NbSTIR1/2* compromised stomatal immunity in *N. benthamiana*. (A) Quantitative real-time PCR (RT-qPCR) analysis of the relative total abundance of *NbSTIR1/2* in *N. benthamiana* (*Nb*) *TRV-GUS* and *TRV-NbSTIR1/2* plants. Data are means ± SD (n= 3). (B) Disease symptoms and bacterial populations in *Nb TRV-GUS* and *TRV-NbSTIR1/2* at 7 d (upper panel), and at 0 or 4 d (lower panel) post spray-inoculation of *Pst* DC3000 ΔHopQ1. Data are means ± SD (n = 4). (C) Stomata apertures of *Nb TRV-GUS* and *TRV-NbSTIR1/2* after 1 h of flood treatment without (mock) or with *Pst* DC3000 ΔHopQ1. Data are means ± SD (n = 50). Scale bars, 5 μm. (D) Bacterial entry assay showing images and quantification of entered bacteria in leaves of *Nb TRV-GUS* and *TRV-NbSTIR1/2.* Data are means ± SD (n = 3). Scale bars, 1 cm. Data were analyzed by one-way ANOVA by Tukey’s post hoc test (P < 0.05; B, C) or two-sided Student’s *t*-test (A, D). n = independent biological replicates (A, B, D).

**Figure S3.**
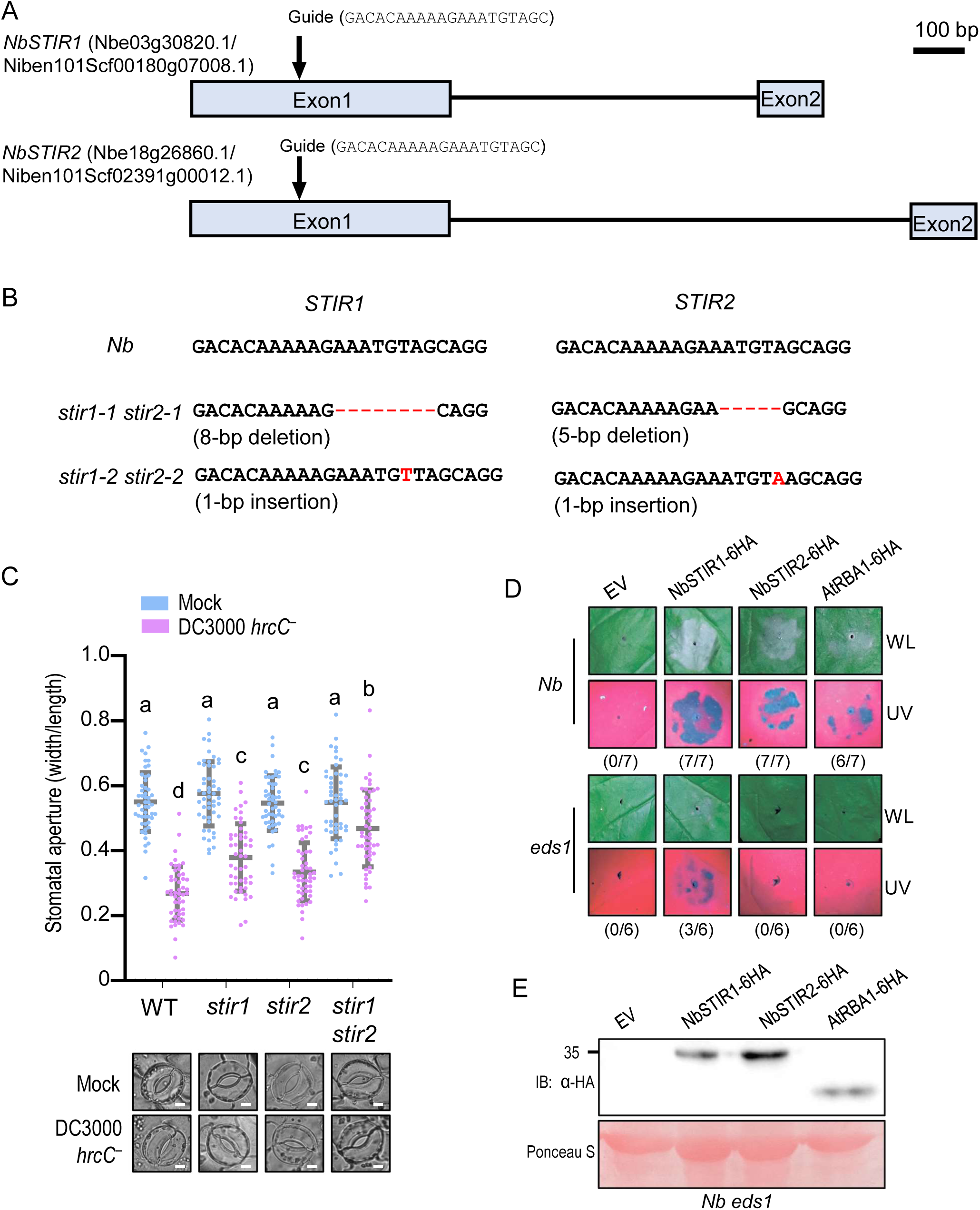
Generation of *stir1 stir2* gene-editing mutants and hypersensitive cell death phenotypes caused by NbSTIR1/2 overexpression. (A) Schematic diagrams showing the guide sequence targeting the first exons of *NbSTIR1* and *NbSTIR2*. (B) The mutations of *NbSTIR1*/2 generated by CRISPR/Cas9 technique in the *stir1-1 stir2-1* and *stir1-2 stir2-2* mutants. (C) Apertures and images of stomata in wild-type *Nb*, *stir1-1*, *stir2-1*, and *stir1-1 stir2-1* mutants after 1 h of flood treatment without (Mock) or with *Pst* DC3000 *hrcC*^−^. Data are means ± SD (n = 50). Scale bars, 5 μm. Data were analyzed by one-way ANOVA by Tukey’s post hoc test (P < 0.05; C). (D-E) Overexpression of NbSTIR1 and NbSTIR2 caused hypersensitive cell death in *Nb* leaves in an *NbEDS1*-dependent manner. Cell death phenotypes (D) were imaged at 2 d after infiltration of *Agrobacterium* strains containing the indicated constructs or empty vector (EV) in wild-type or *eds1* under white light (WL) or UV light (UV). The numbers in parentheses represent the proportion of leaves exhibiting cell death out of the total number of infiltrated leaves. Protein levels in *Nb eds1* leaves were detected by immunoblotting with anti-HA antibody (E). Ponceau-S staining of Rubisco was used as a loading control.

**Figure S4.**
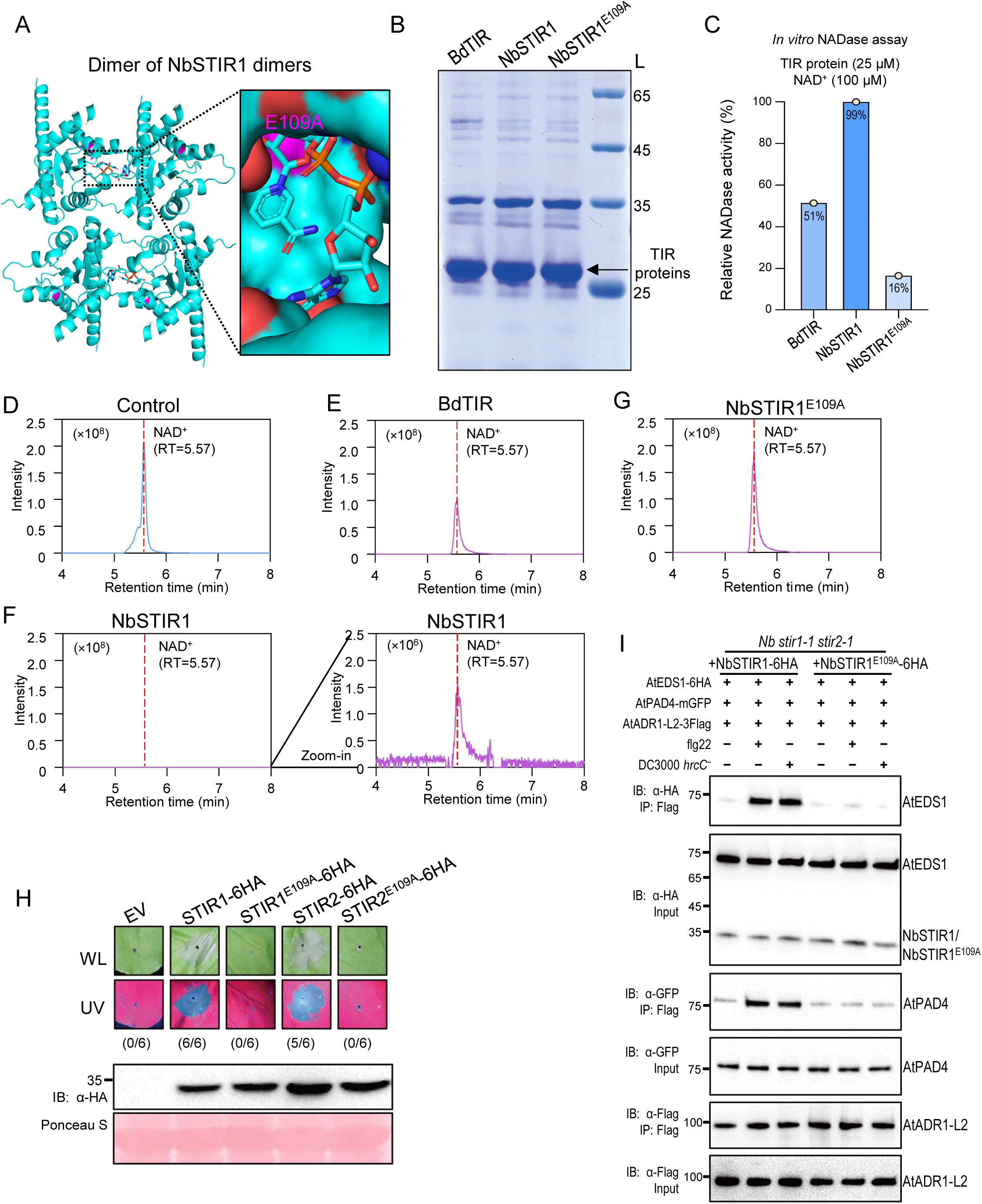
NbSTIR1 is an NAD^+^-cleaving enzyme that relies on the conserved glutamate E109. (A) A schematic diagram of overall structure of dimer of NbSTIR1 dimers predicted by AlphaFold 3. Catalytic centers involve the BB-loops and form in the head-to-tail TIR dimers. The conserved E residue within the catalytic center of the head-to-tail TIR dimers for the NADase activity is marked. (B) SDS-PAGE analysis showing the purified TIR proteins used for the NADase activity assay. His-tagged BdTIR, NbSTIR1, NbSTIR1^E109A^ were purified from *E*. *coli*. Arrowheads indicate the expected protein size. L, protein ladder. (C) *In vitro* NADase activity of BdTIR, NbSTIR1 or NbSTIR1^E109A^ via UPLC-MS/MS. Reaction completion (%) of each sample was utilized to represent the relative NADase activity (%). (D-G) Ultra performance liquid chromatography (UPLC) analysis of NAD^+^ following a NADase activity assay without (control, D) or with BdTIR (E), NbSTIR1 (F), or NbSTIR1^E109A^ (G). (H) Hypersensitive cell death phenotypes at 2 day after infiltration of *Agrobacterium* strains containing the indicated vectors including empty vector (EV), NbSTIR1-6HA, NbSTIR1^E109A^-6HA, NbSTIR2-6HA, or NbSTIR2^E109A^-6HA in *Nb* leaves, and immunoblotting analysis of their protein levels. The numbers in parentheses represent the proportion of leaves exhibiting cell death out of the total number of infiltrated leaves. Ponceau-S staining of Rubisco was used as a loading control. (I) Co-IP assay showed that complementing *NbSTIR1* expression restored *Pst* DC3000 *hrcC*^−^ and flg22-induced association of AtADR1-L2 with AtEDS1 and AtPAD4 in *stir1 stir2* leaves, but complementing NbSTIR1^E109A^ expression did not. The total proteins were immunoprecipitated with the anti-GFP agarose beads, and the IP products were detected by immunoblotting using anti-GFP or anti-HA antibody. IB, immunoblotting; IP, immunoprecipitation.

**Figure S5.**
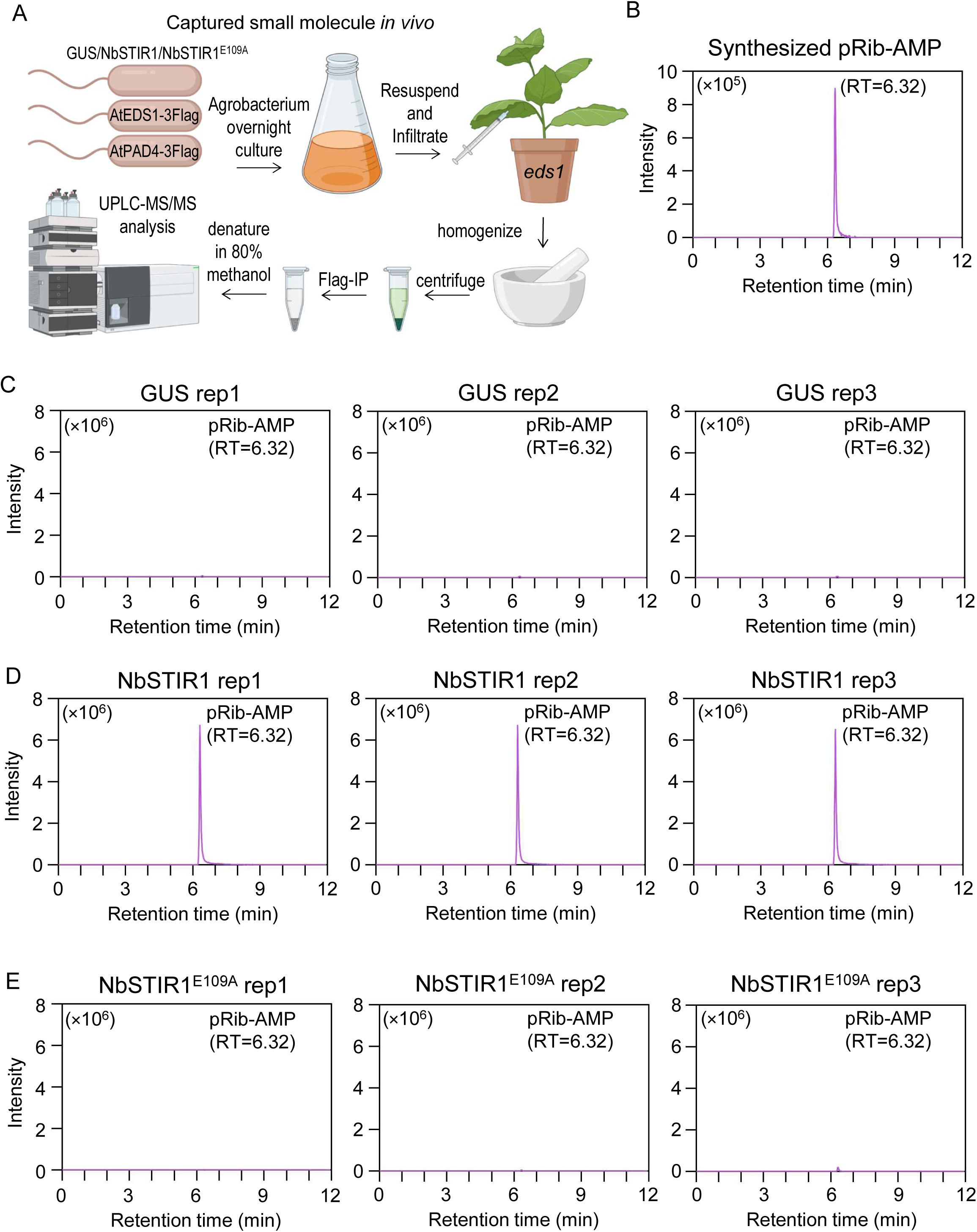
NbSTIR1 but not NbSTIR1^E109A^ produces the immune molecule pRib-AMP in *N. benthamiana.* (A) Flowchart for analyses of pRib-AMP in *Nb* leaves. AtEDS1-3Flag and AtPAD4-3Flag were coexpressed with GUS, NbSTIR1, or NbSTIR1^E109A^ in *Nb eds1* leaves, and the AtEDS1-AtPAD4 complex was purified by anti-Flag affinity chromatography, and pRib-AMP captured in the purified AtEDS1-AtPAD4 complex was analyzed by UPLC-MS/MS. (B-E) UPLC-MS/MS analysis of the synthesized standard pRib-AMP (B), and the AtEDS1-AtPAD4-bound pRib-AMP generated by coexpression with GUS (C), NbSTIR1 (D), or NbSTIR1^E109A^ (E).

**Figure S6.**
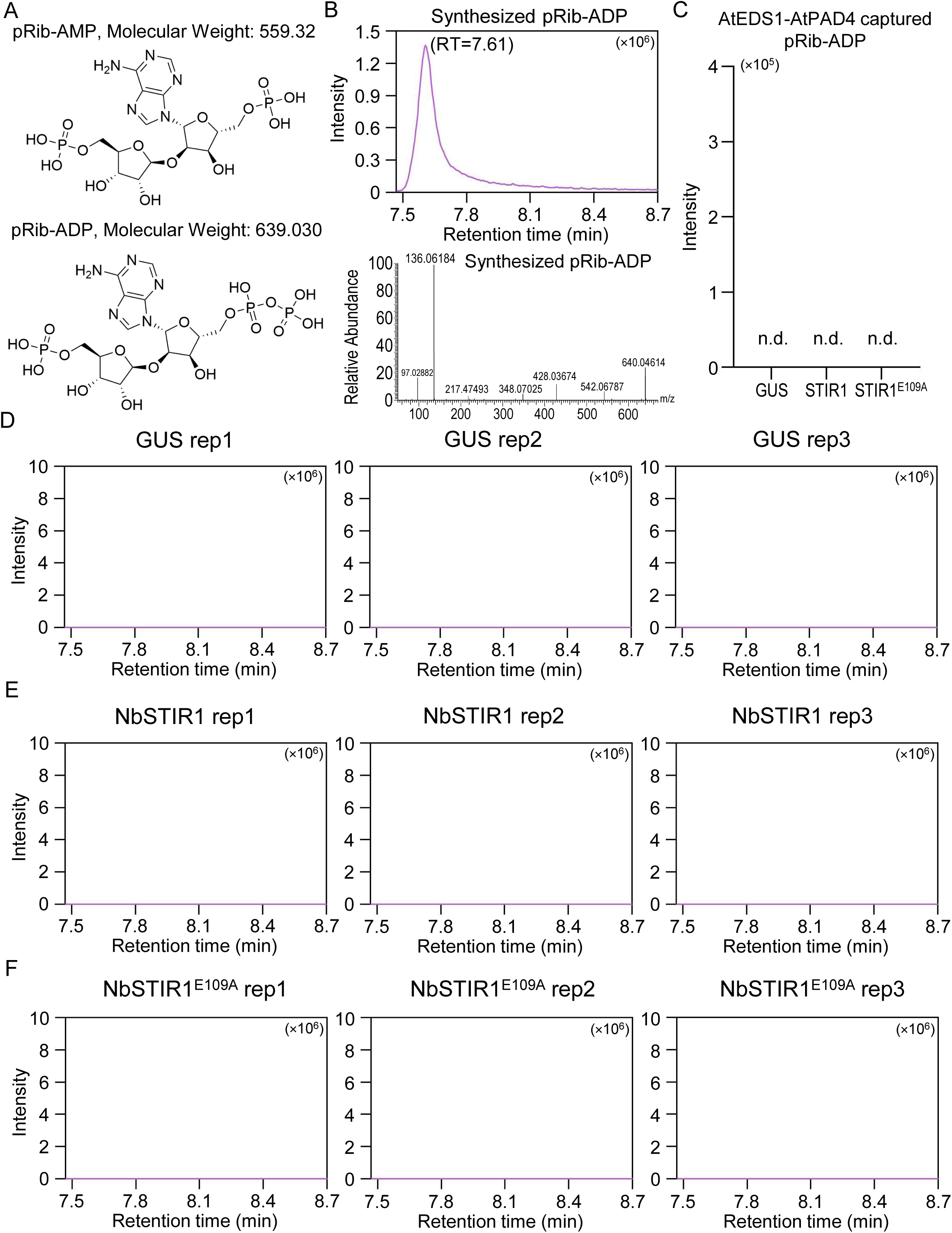
pRib-ADP is not detected within the AtEDS1-AtPAD4 complex purified from *N. benthamiana*. (A) Structure structure and theoretical molecular weight of pRib-AMP (upper panel) and pRib-ADP (lower panel). (B) UPLC-MS/MS analysis of the synthesized standard pRib-ADP. (C-F) UPLC-MS/MS analysis of pRib-ADP within AtEDS1-AtPAD4 complex (C) coexpressed with GUS (D), NbSTIR1 (E), or NbSTIR1^E109A^ (F). n.d., not detected.

**Figure S7.**
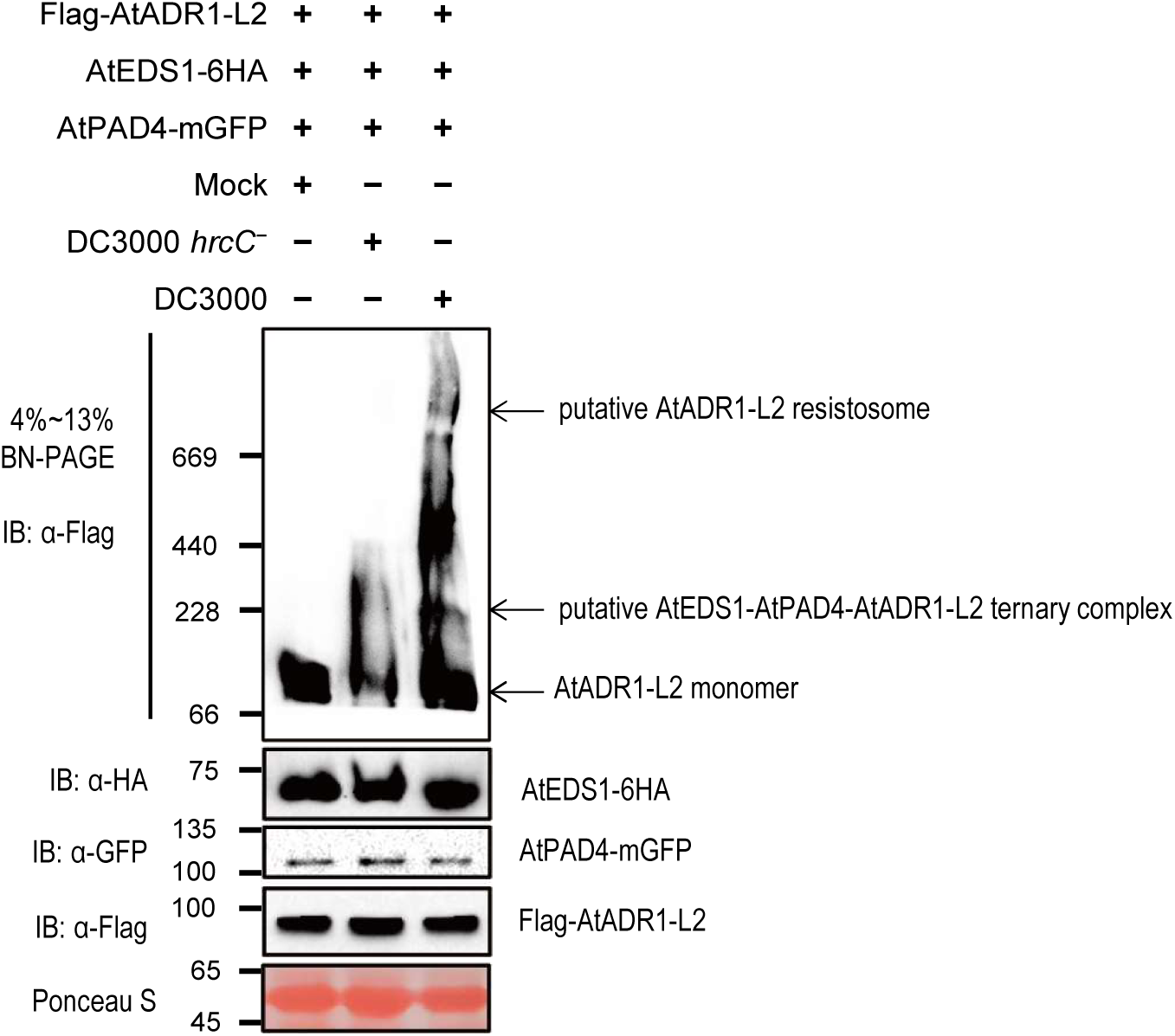
The oligomeric AtADR1-L2 resistosome is not induced upon *Pst* DC3000 *hrcC*^−^ infection. BN-PAGE analysis revealed that *Pst* DC3000 *hrcC*^−^ infection did not promote the formation of oligomeric AtADR1-L2 resistosome in *Nb eds1* leaves, but triggered a lower molecular weight complex containing AtADR1-L2, which was putatively the AtEDS1-AtPAD4-AtADR1- L2 ternary complex. Ponceau-S staining of Rubisco was used as a loading control. IB, immunoblotting.

**Figure S8.**
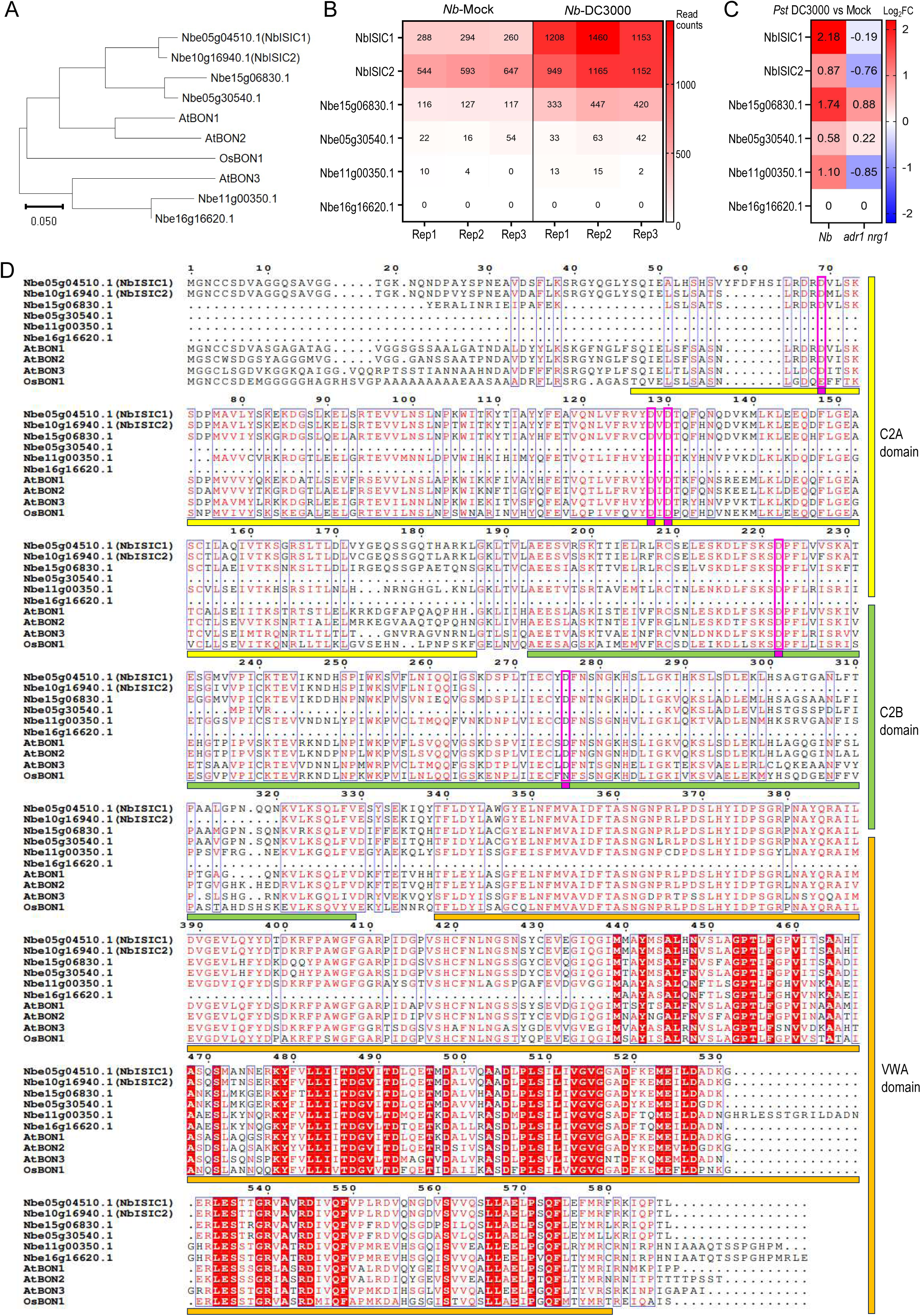
Phylogenetic analysis and protein alignment of NbISIC1/2 and homologs. (A) Phylogenetic analysis of two C2-domain proteins NbISIC1, NbISIC2, their homologs from *Nb*, *Arabidopsis* (AtBON1/AT5G61900, AtBON2/AT5G07300, AtBON3/AT1G08860), Oryza sativa (OsBON1, Os02g0521300). (B-C) Expression levels (B) and fold changes (C) of *NbISIC1*, *NbISIC2*, and their *Nb* homolog genes from RNA-seq analysis of *Nb* wildtype and *adr1 nrg1* leaves under mock or *Pst* DC3000 treatment. (D) Multiple protein sequence alignment of NbISIC1, NbISIC2, their Nb homologs, AtBON1, AtBON2, AtBON3, and OsBON1. The conserved C2 domains, VWA domains, and conserved D residues within C2 domains for Ca^2+^ binding are marked.

**Figure S9.**
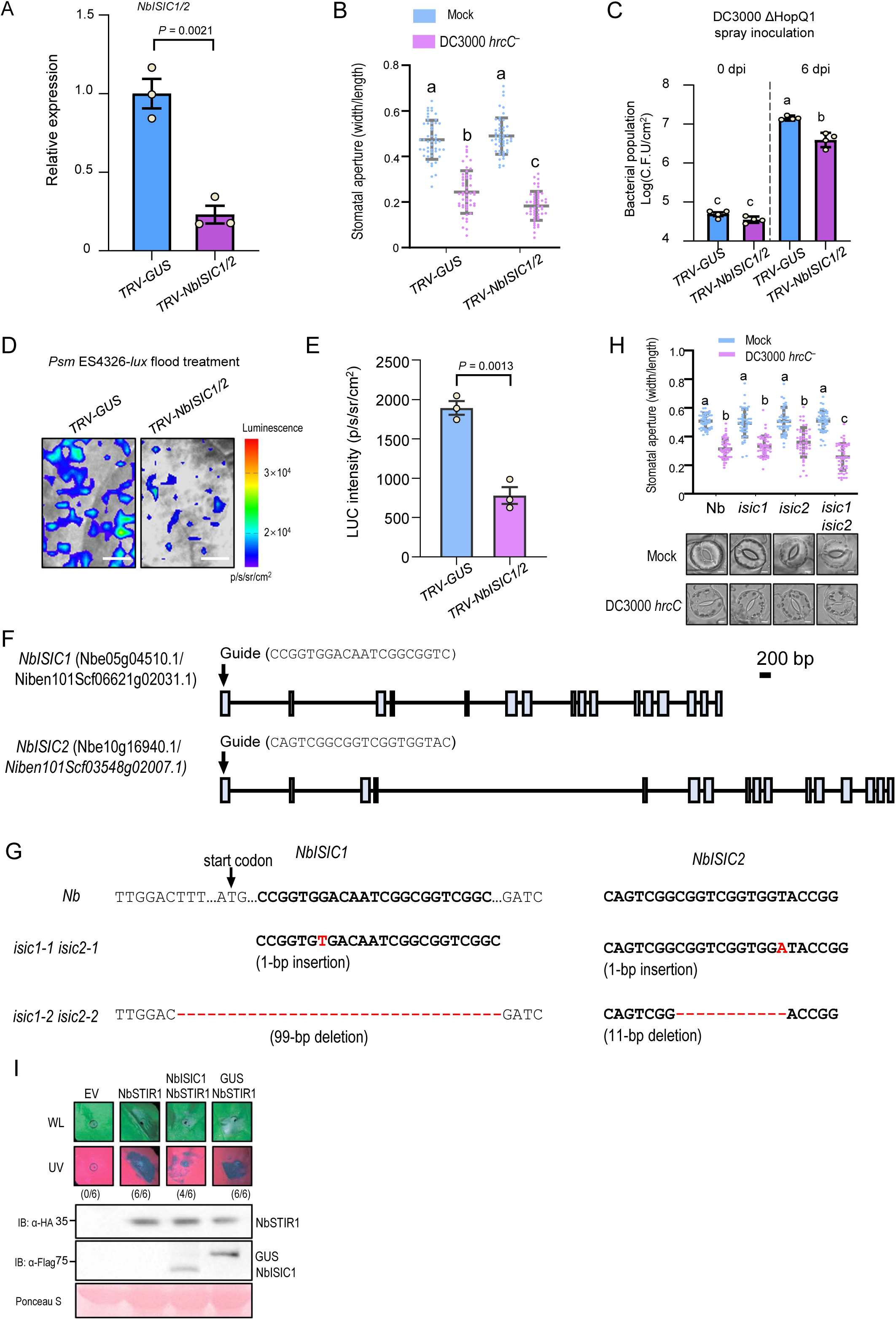
NbISIC1/2 regulates stomatal immunity. (A) RT-qPCR analysis of the relative total abundance of *NbISIC1/2* in *Nb TRV-GUS* and *TRV-NbISIC1/2* plants. Data are means ±SD (n = 3). (B) Stomatal apertures of *TRV-GUS* or *TRV-NbISIC1/2* after 1 h of flood treatment with mock or *Pst* DC3000 *hrcC*^−^. Data are means ± SD (n = 50). (C) Bacterial populations in leaves of *TRV-GUS* or *TRV-NbISIC1/2* at 0 or 6 d post spray-inoculation of *Pst* DC3000 ΔHopQ1. Data are means ± SD (n = 4). (D-E) Bacterial entry assay showing images (D) and quantification (E) of bacteria in leaves of *TRV-GUS* or *TRV-NbISIC1/2.* Data are means ± SD (n = 3). Scale bars, 0.5 cm. (F) Schematic diagrams showing the guide sequence targeting the first exons of *NbISIC1/2*. (G) The mutations of *NbISIC1*/2 generated by CRISPR/Cas9 technique in the *isic1-1 isic2-1* and *isic1-2 isic2-2* mutants. (H) Apertures and images of stomata in wild-type *Nb*, *isic1-1*, *isic2-1*, and *isic1-1 isic2-1* mutants after 1 h of flood treatment without (Mock) or with *Pst* DC3000 *hrcC*^−^. Data are means ± SD (n = 50). Scale bars, 5 μm. (I) NbISIC1 overexpression attenuated NbSTIR1-induced hypersensitive cell death in *Nb*leaves. Images were captured at 2 d after infiltration of *Agrobacterium* strains containing the indicated constructs or empty vector (EV) in *Nb* leaves (upper panel), and proteins were detected by anti-HA or anti-Flag antibody (lower panel). The numbers in parentheses represent the proportion of leaves exhibiting cell death out of the total number of infiltrated leaves. Ponceau-S staining of Rubisco served as a loading control. Data were analyzed by one-way ANOVA by Tukey’s post hoc test (P < 0.05; B, C, H) or two-sided Student’s *t*-test (A, E); n = independent biological replicates (A, C, E).

**Figure S10.**
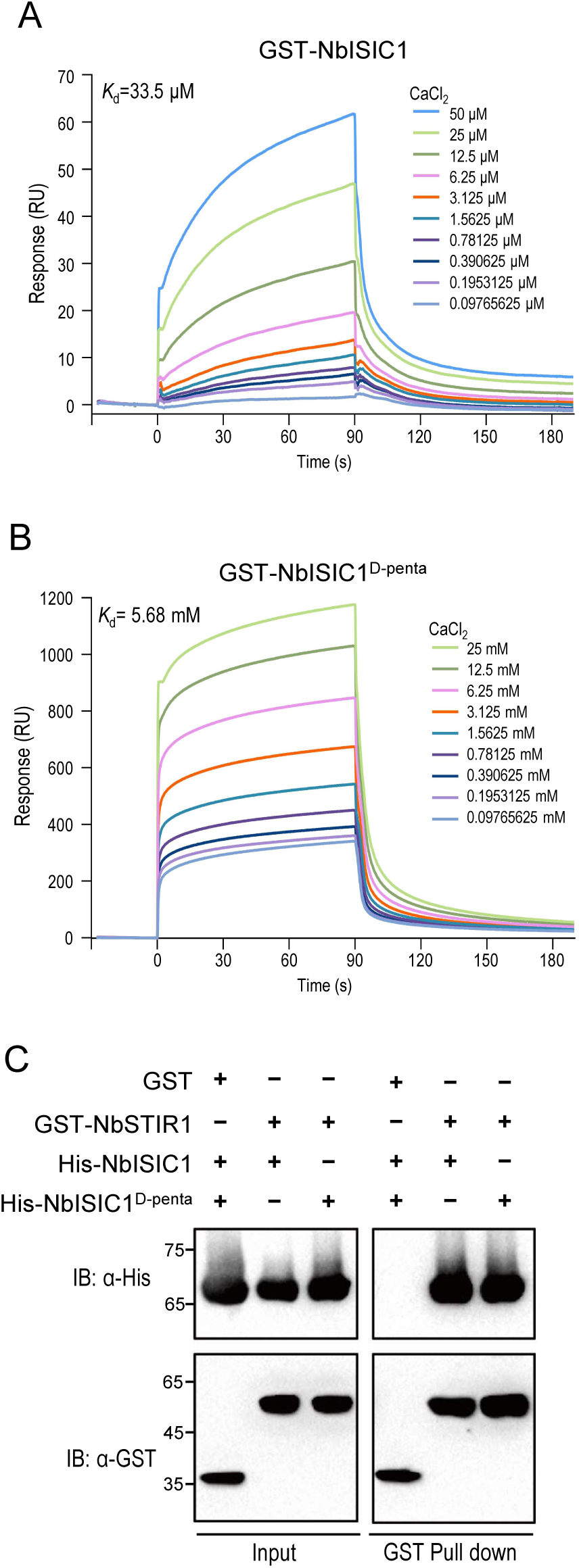
NbISIC1 binds to Ca^2+^ and interacts with NbSTIR1. (A-B) Surface plasmon resonance analysis of NbISIC1 (A) and NbISIC1^D-penta^ (D69/128/130/222/276N) (B) binding to Ca^2+^. *K*_d_, dissociation constant. (C) Pull-down assays show that both NbISIC1 and NbISIC1^D-penta^ interact with NbSTIR1 *in vitro*. Pull-down assays were performed as in Figure 4J.

**Figure S11.**
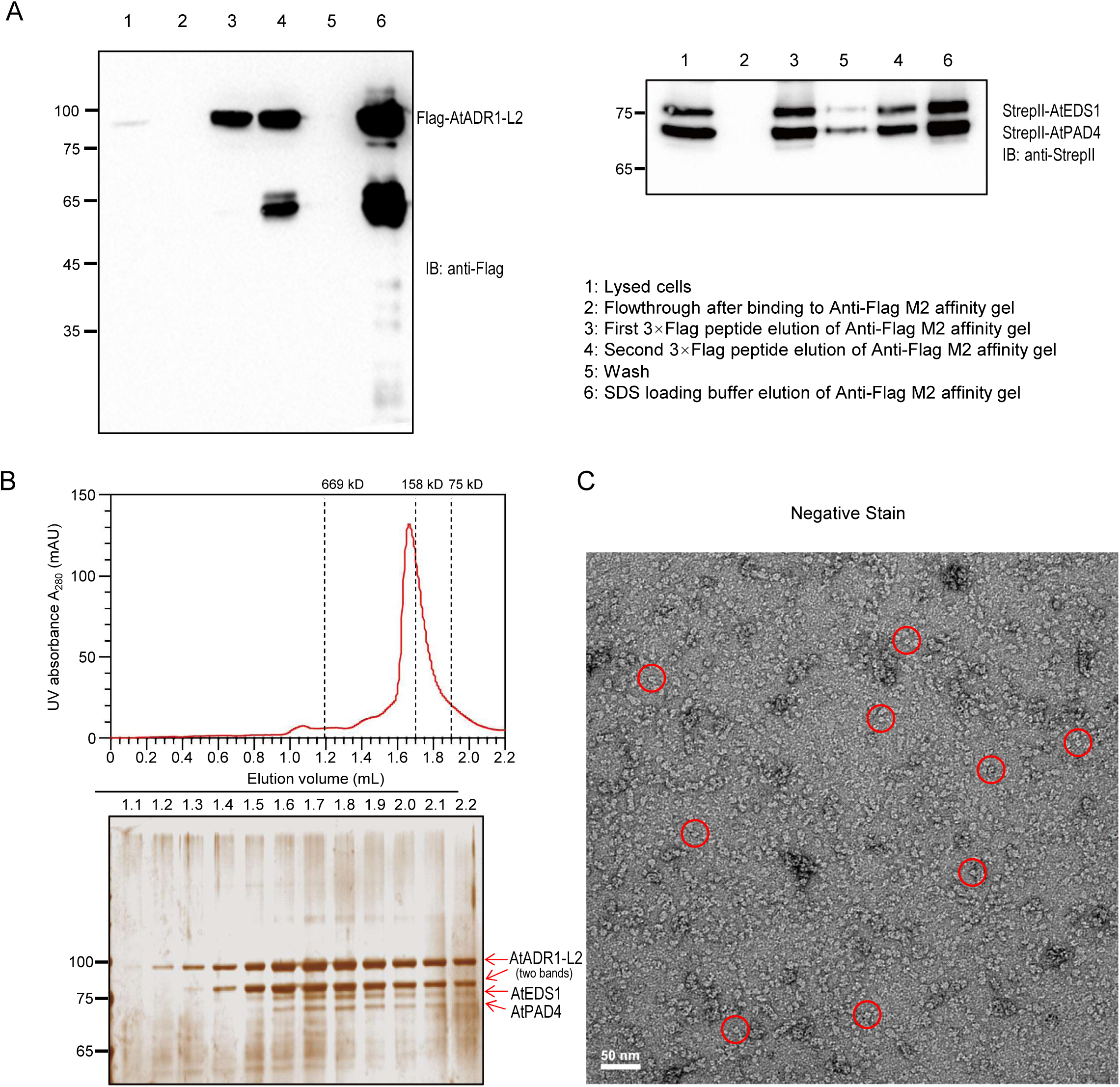
Purification of the AtEDS1-AtPAD4-AtADR1-L2 complex expressed in *N. benthamiana* leaves. (A) Anti-Flag affinity chromotography of the AtEDS1-AtPAD4-AtADR1-L2 complex, with samples analyzed by immunoblot. StrepII-AtEDS1, StrepII-AtPAD4 and Flag-AtADR1-L2 were coexpressed in *Nb eds1* leaves with *Pst* DC3000 *hrcC*^−^ infection. (B) Gel filtration of the AtEDS1-AtPAD4-AtADR1-L2 complex. The fractions were analyzed by silver-staining. Gel filtration molecular weight markers were indicated by dash lines. (C) Representative negative staining image of the AtEDS1-AtPAD4-AtADR1-L2 complex, with particles indicated by red circles.

**Figure S12.**
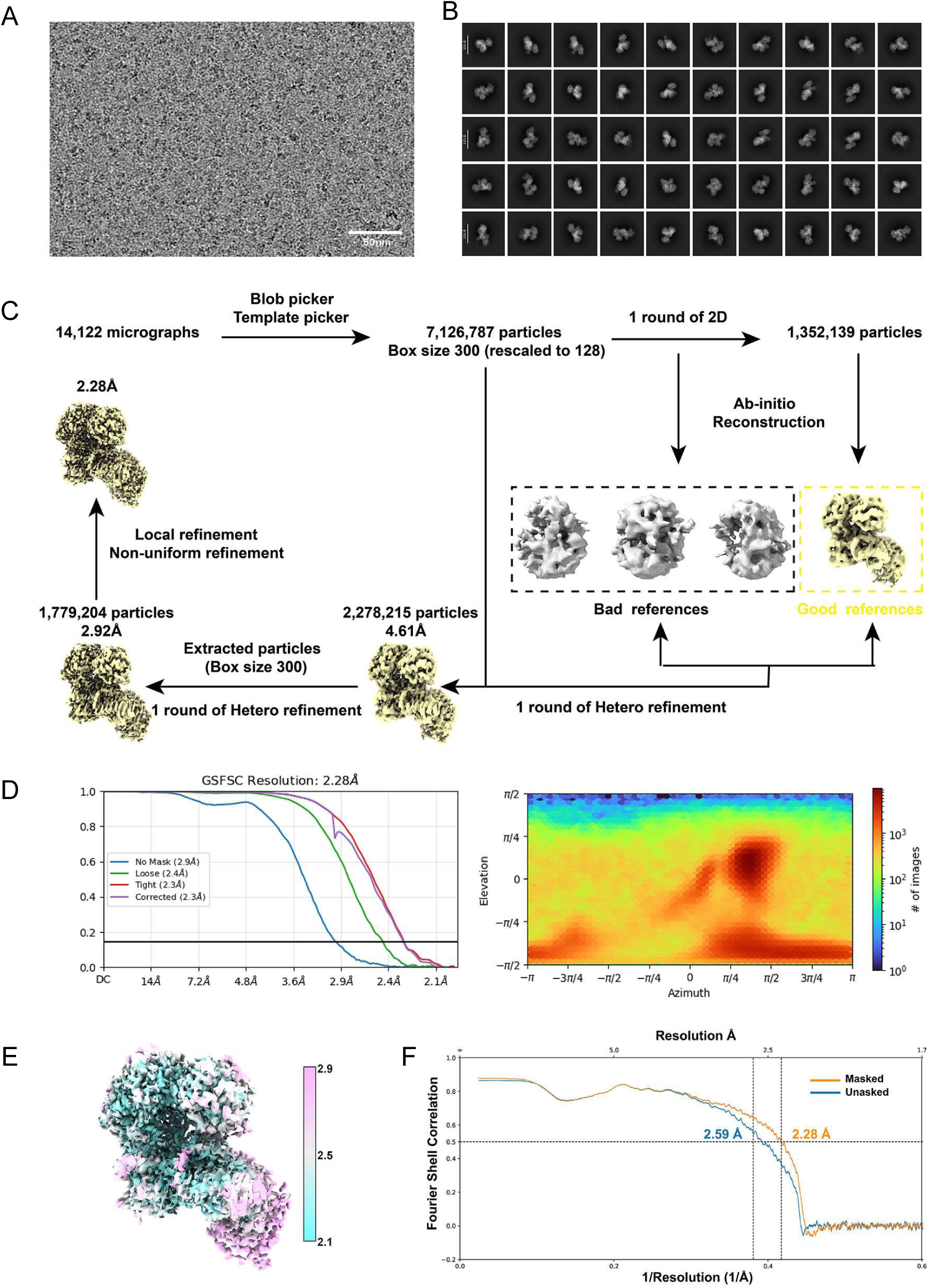
3D reconstruction of the AtEDS1-AtPAD4-AtADR1-L2 complex. (A-B) Representative cryo-EM micrograph (A) and 2D class averages (B) of AtEDS1-AtPAD4- AtADR1-L2 complex. (C) Flowchart and data processing for AtEDS1-AtPAD4-AtADR1-L2 3D reconstruction. (D) Gold standard Fourier shell correlation (GSFSC) resolution plot and orientation distribution of AtEDS1-AtPAD4-AtADR1-L2 map. (E-F) The final EM density map (E) and FSC curves (F) of the AtEDS1-AtPAD4-AtADR1-L2 complex with the local resolution estimated by ResMap.

**Figure S13.**
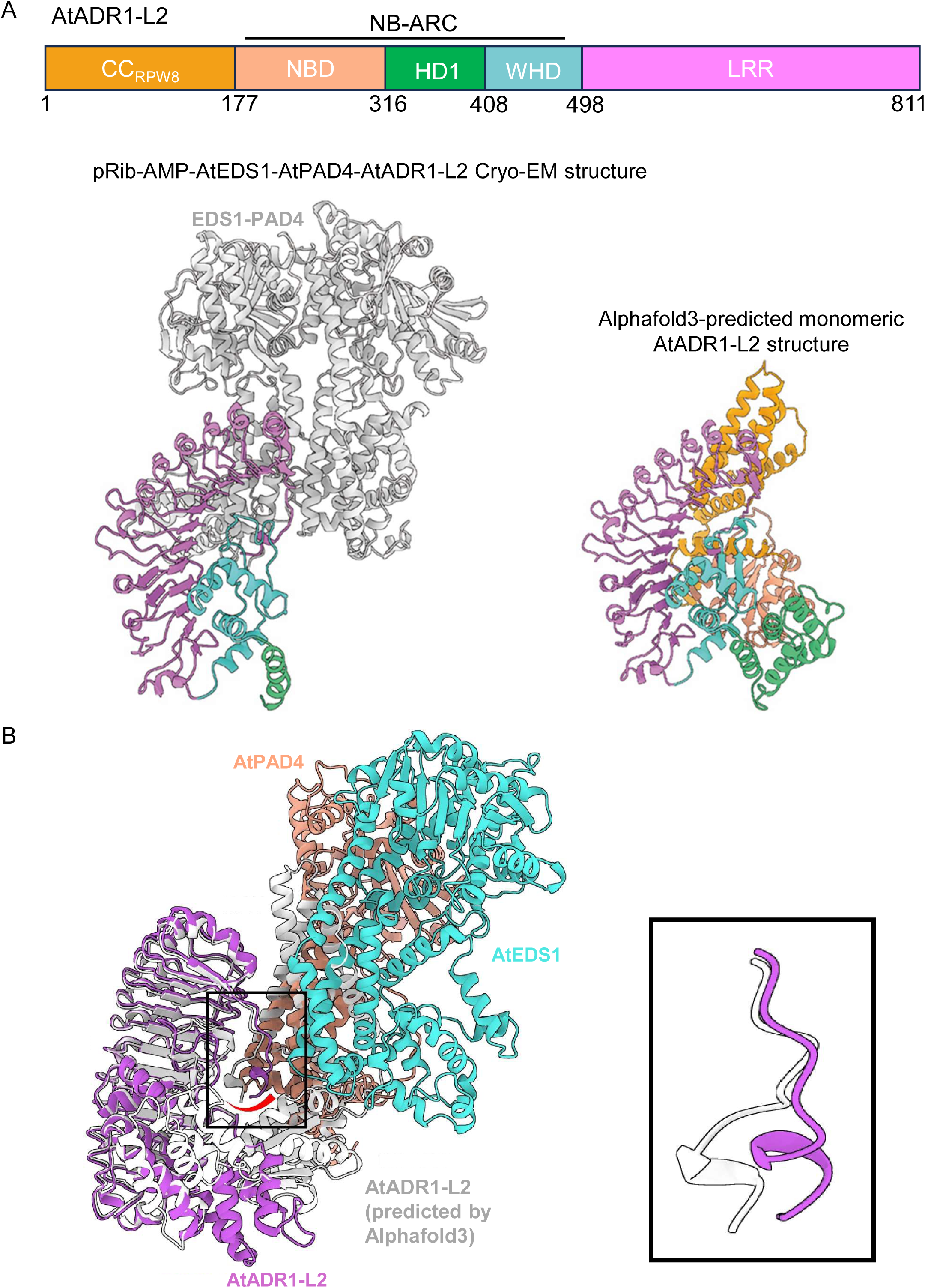
Structural alignment of AtEDS1-AtPAD4-AtADR1-L2 and the predicted AtADR1-L2 monomeric structure. (A) Schematic diagram of AtADR1-L2 domains (upper panel), and structural comparison of the pRib-AMP-bound-AtEDS1-AtPAD4-AtADR1-L2 structure (bottom left) and the monomeric ADR1-L2 structure predicted by Alphafold3 (bottom right). AtEDS1-AtPAD4 is shown in gray. For AtADR1-L2, the CC_RPW8_, NBD, HD1, WHD and LRR domains are shown in orange, beige, green, cyan, and pink respectively. (B) Structural alignment of the pRib-AMP-bound-AtEDS1-AtPAD4-AtADR1-L2 structure and the monomeric AtADR1-L2 structure predicted by Alphafold3 shown in gray (left) with the C-terminal loop of AtADR1-L2 enlarged (right). Color codes for AtEDS1, AtPAD4, and AtADR1- L2 are the same as in Figure 5-6. The monomeric AtADR1-L2 structure predicted by Alphafold3 is shown in gray.

**Figure S14.**
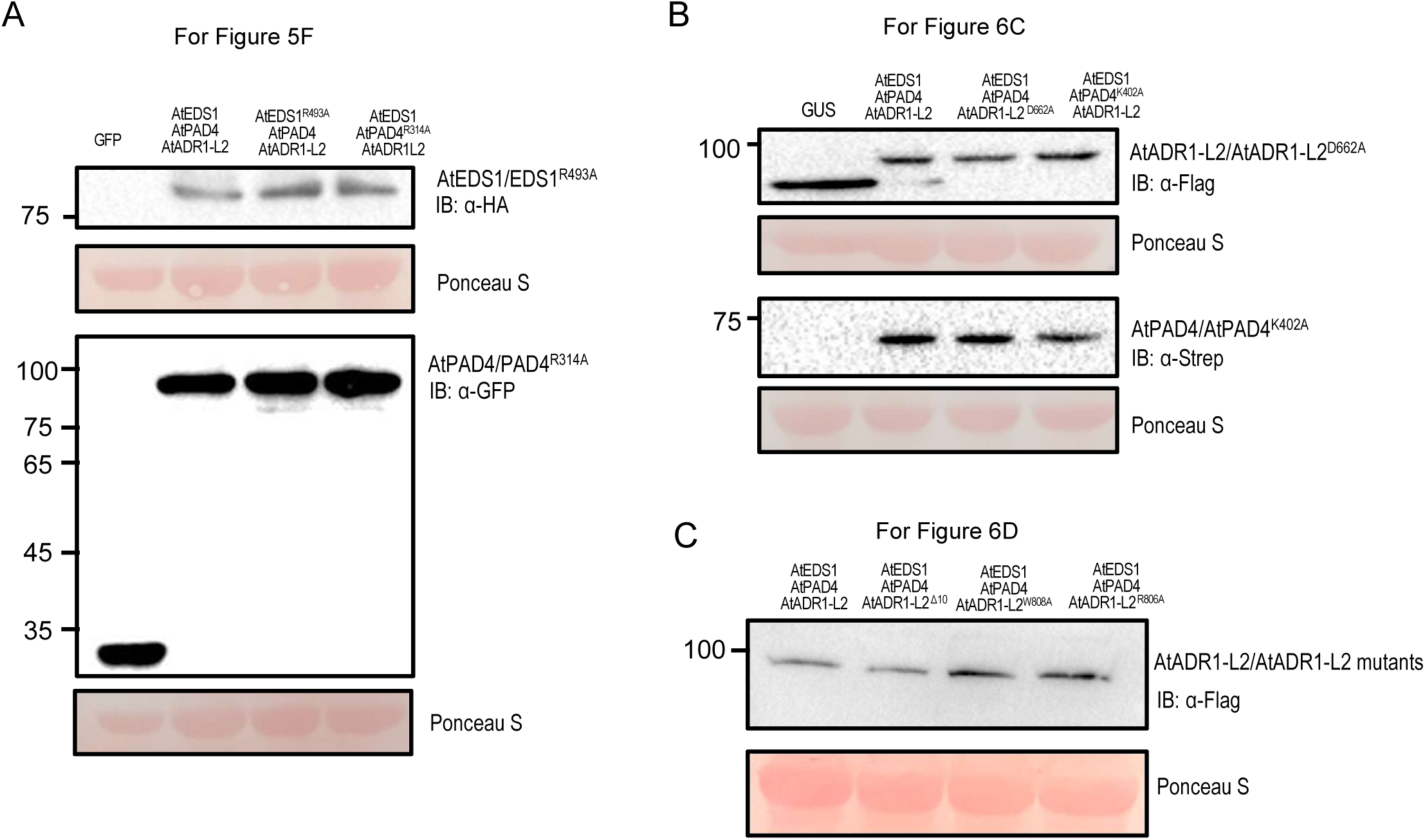
Protein expression of AtEDS1, AtPAD4, AtADR1-L2, and their variants. (A-C) Protein expression of AtEDS1, AtPAD4, AtADR1-L2, and their variants in *Nb eds1* leaves as in Figures 5F (A), 6C (B), and 6D (C) was detected by immunoblotting with anti-HA, anti-GFP, anti-Flag, or anti-Strep antibody. Ponceau-S staining of Rubisco was used as a loading control.

Table S1. Cryo-EM data collection, refinement and validation statistics.

Table S2. Primer sequences used in this study

## Notes

### Competing Interest Statement

The authors have declared no competing interest.

## References

Abramson, J., Adler, J., Dunger, J., Evans, R., Green, T., Pritzel, A., Ronneberger, O., Willmore, L., Ballard, A.J., Bambrick, J., Bodenstein, S.W., Evans, D.A., Hung, C.-C., O’Neill, M., Reiman, D., Tunyasuvunakool, K., Wu, Z., Žemgulytė, A., Arvaniti, E., Beattie, C., Bertolli, O., Bridgland, A., Cherepanov, A., Congreve, M., Cowen-Rivers, A.I., Cowie, A., Figurnov, M., Fuchs, F.B., Gladman, H., Jain, R., Khan, Y.A., Low, C.M.R., Perlin, K., Potapenko, A., Savy, P., Singh, S., Stecula, A., Thillaisundaram, A., Tong, C., Yakneen, S., Zhong, E.D., Zielinski, M., Žídek, A., Bapst, V., Kohli, P., Jaderberg, M., Hassabis, D., and Jumper, J.M. (2024). Accurate structure prediction of biomolecular interactions with AlphaFold 3. Nature 630, 493–500.

Adachi, H., Derevnina, L., and Kamoun, S. (2019). NLR singletons, pairs, and networks: evolution, assembly, and regulation of the intracellular immunoreceptor circuitry of plants. Curr Opin Plant Biol 50, 121–131.

Afonine, P.V., Poon, B.K., Read, R.J., Sobolev, O.V., Terwilliger, T.C., Urzhumtsev, A., and Adams, P.D. (2018). Real-space refinement in PHENIX for cryo-EM and crystallography. Acta Crystallographica Section D: Structural Biology 74, 531–544.

Amunts, A., Brown, A., Bai, X.-c., Llácer, J.L., Hussain, T., Emsley, P., Long, F., Murshudov, G., Scheres, S.H., and Ramakrishnan, V. (2014). Structure of the yeast mitochondrial large ribosomal subunit. Science 343, 1485–1489.

Bi, G.Z., Su, M., Li, N., Liang, Y., Dang, S., Xu, J.C., Hu, M.J., Wang, J.Z., Zou, M.X., Deng, Y.A., Li, Q.Y., Huang, S.J., Li, J.J., Chai, J.J., He, K.M., Chen, Y.H., and Zhou, J.M. (2021). The ZAR1 resistosome is a calcium-permeable channel triggering plant immune signaling. Cell 184, 3528–3541.

Collier, S.M., Hamel, L.P., and Moffett, P. (2011). Cell death mediated by the N-terminal domains of a unique and highly conserved class of NB-LRR protein. Mol Plant Microbe Interact 24, 918–931.

Couto, D., and Zipfel, C. (2016). Regulation of pattern recognition receptor signalling in plants. Nat Rev Immunol 16, 537–552.

Cui, H., Tsuda, K., and Parker, J.E. (2015). Effector-triggered immunity: from pathogen perception to robust defense. Annu Rev Plant Biol 66, 487–511.

Dharmasiri, N., Dharmasiri, S., and Estelle, M. (2005). The F-box protein TIR1 is an auxin receptor. Nature 435, 441–445.

Emsley, P., and Cowtan, K. (2004). Coot: model-building tools for molecular graphics. Acta crystallographica section D: biological crystallography 60, 2126–2132.

Feehan, J.M., Castel, B., Bentham, A.R., and Jones, J.D. (2020). Plant NLRs get by with a little help from their friends. Curr Opin Plant Biol 56, 99–108.

Ge, Z., Bergonci, T., Zhao, Y., Zou, Y., Du, S., Liu, M.C., Luo, X., Ruan, H., Garcia-Valencia, L.E., Zhong, S., Hou, S., Huang, Q., Lai, L., Moura, D.S., Gu, H., Dong, J., Wu, H.M., Dresselhaus, T., Xiao, J., Cheung, A.Y., and Qu, L.J. (2017). Arabidopsis pollen tube integrity and sperm release are regulated by RALF-mediated signaling. Science 358, 1596–1600.

Gong, Y., Tian, L., Kontos, I., Li, J., and Li, X. (2023). Plant immune signaling network mediated by helper NLRs. Curr Opin Plant Biol 73, 102354.

Grant, J.J., Chini, A., Basu, D., and Loake, G.J. (2003). Targeted activation tagging of the Arabidopsis NBS-LRR gene, ADR1, conveys resistance to virulent pathogens. Mol Plant Microbe Interact 16, 669–680.

Horsefield, S., Burdett, H., Zhang, X., Manik, M.K., Shi, Y., Chen, J., Qi, T., Gilley, J., Lai, J.S., Rank, M.X., Casey, L.W., Gu, W., Ericsson, D.J., Foley, G., Hughes, R.O., Bosanac, T., von Itzstein, M., Rathjen, J.P., Nanson, J.D., Boden, M., Dry, I.B., Williams, S.J., Staskawicz, B.J., Coleman, M.P., Ve, T., Dodds, P.N., and Kobe, B. (2019). NAD(+) cleavage activity by animal and plant TIR domains in cell death pathways. Science 365, 793–799.

Hou, S., Rodrigues, O., Liu, Z., Shan, L., and He, P. (2024). Small holes, big impact: stomata in plant-pathogen-climate epic trifecta. Mol Plant 17, 26–49.

Huang, S., Jia, A., Song, W., Hessler, G., Meng, Y., Sun, Y., Xu, L., Laessle, H., Jirschitzka, J., Ma, S., Xiao, Y., Yu, D., Hou, J., Liu, R., Sun, H., Liu, X., Han, Z., Chang, J., Parker, J.E., and Chai, J. (2022). Identification and receptor mechanism of TIR-catalyzed small molecules in plant immunity. Science 377, eabq3297.

Hussain, M.Z., Ghani, Q.P., and Hunt, T.K. (1989). Inhibition of prolyl hydroxylase by poly(ADP-ribose) and phosphoribosyl-AMP: Possible role of ADP-ribosylation in intracellular prolyl hydroxylase regulation. Journal of Biological Chemistry 264, 7850–7855.

Jacob, P., Kim, N.H., Wu, F., El-Kasmi, F., Chi, Y., Walton, W.G., Furzer, O.J., Lietzan, A.D., Sunil, S., Kempthorn, K., Redinbo, M.R., Pei, Z.M., Wan, L., and Dangl, J.L. (2021). Plant “helper” immune receptors are Ca(2+)-permeable nonselective cation channels. Science 373, 420–425.

Jacob, P., Hige, J., Song, L., Bayless, A., Russ, D., Bonardi, V., El Kasmi, F., Wunsch, L., Yang, Y., Fitzpatrick, C.R., McKinney, B.J., Nishimura, M.T., Grant, M.R., and Dangl, J.L. (2023). Broader functions of TIR domains in Arabidopsis immunity. Proc Natl Acad Sci U S A 120, e2220921120.

Jia, A., Huang, S., Song, W., Wang, J., Meng, Y., Sun, Y., Xu, L., Laessle, H., Jirschitzka, J., Hou, J., Zhang, T., Yu, W., Hessler, G., Li, E., Ma, S., Yu, D., Gebauer, J., Baumann, U., Liu, X., Han, Z., Chang, J., Parker, J.E., and Chai, J. (2022). TIR-catalyzed ADP-ribosylation reactions produce signaling molecules for plant immunity. Science 377, eabq8180.

Johanndrees, O., Baggs, E.L., Uhlmann, C., Locci, F., Lässle, H.L., Melkonian, K., Käufer, K., Dongus, J.A., Nakagami, H., Krasileva, K., Parker, J.E., and Lapin, D. (2023). Variation in plant Toll/Interleukin-1 receptor domain protein dependence on ENHANCED DISEASE SUSCEPTIBILITY 1. Plant Physiol 191, 626–642.

Jones, J.D., and Dangl, J.L. (2006). The plant immune system. Nature 444, 323–329.

Jones, J.D., Vance, R.E., and Dangl, J.L. (2016). Intracellular innate immune surveillance devices in plants and animals. Science 354.

Jubic, L.M., Saile, S., Furzer, O.J., El Kasmi, F., and Dangl, J.L. (2019). Help wanted: helper NLRs and plant immune responses. Curr Opin Plant Biol 50, 82–94.

Koster, P., DeFalco, T.A., and Zipfel, C. (2022). Ca(2+) signals in plant immunity. EMBO J 41, e110741.

Lapin, D., Johanndrees, O., Wu, Z., Li, X., and Parker, J.E. (2022). Molecular innovations in plant TIR-based immunity signaling. Plant Cell 34, 1479–1496.

Liang, X., and Zhou, J.M. (2018). Receptor-like cytoplasmic kinases: central players in plant receptor kinase-mediated signaling. Annu Rev Plant Biol 69, 267–299.

Liebschner, D., Afonine, P.V., Baker, M.L., Bunkóczi, G., Chen, V.B., Croll, T.I., Hintze, B., Hung, L.-W., Jain, S., and McCoy, A.J. (2019). Macromolecular structure determination using X-rays, neutrons and electrons: recent developments in Phenix. Acta Crystallographica Section D: Structural Biology 75, 861–877.

Liu, Y., Schiff, M., Marathe, R., and Dinesh-Kumar, S.P. (2002). Tobacco Rar1, EDS1 and NPR1/NIM1 like genes are required for N-mediated resistance to tobacco mosaic virus. Plant J 30, 415–429.

Liu, Z., Hou, S., Rodrigues, O., Wang, P., Luo, D., Munemasa, S., Lei, J., Liu, J., Ortiz-Morea, F.A., Wang, X., Nomura, K., Yin, C., Wang, H., Zhang, W., Zhu-Salzman, K., He, S.Y., He, P., and Shan, L. (2022). Phytocytokine signalling reopens stomata in plant immunity and water loss. Nature 605, 332–339.

Ma, S., Lapin, D., Liu, L., Sun, Y., Song, W., Zhang, X., Logemann, E., Yu, D., Wang, J., Jirschitzka, J., Han, Z., Schulze-Lefert, P., Parker, J.E., and Chai, J. (2020). Direct pathogen-induced assembly of an NLR immune receptor complex to form a holoenzyme. Science 370, eabe3069.

Martin, R., Qi, T., Zhang, H., Liu, F., King, M., Toth, C., Nogales, E., and Staskawicz, B.J. (2020). Structure of the activated ROQ1 resistosome directly recognizing the pathogen effector XopQ. Science 370, eabd9993.

Maruta, N., Sorbello, M., Lim, B.Y.J., McGuinness, H.Y., Shi, Y., Ve, T., and Kobe, B. (2023). TIR domain-associated nucleotides with functions in plant immunity and beyond. Curr Opin Plant Biol 73, 102364.

Melotto, M., Underwood, W., and He, S.Y. (2008). Role of stomata in plant innate immunity and foliar bacterial diseases. Annu Rev Phytopathol 46, 101–122.

Melotto, M., Fochs, B., Jaramillo, Z., and Rodrigues, O. (2024). Fighting for survival at the stomatal gate. Annu Rev Plant Biol 75, 551–577.

Melotto, M., Underwood, W., Koczan, J., Nomura, K., and He, S.Y. (2006). Plant stomata function in innate immunity against bacterial invasion. Cell 126, 969–980.

Ngou, B.P.M., Ahn, H.K., Ding, P., and Jones, J.D.G. (2021). Mutual potentiation of plant immunity by cell-surface and intracellular receptors. Nature 592, 110–115.

Nishimura, M.T., Anderson, R.G., Cherkis, K.A., Law, T.F., Liu, Q.L., Machius, M., Nimchuk, Z.L., Yang, L., Chung, E.H., El Kasmi, F., Hyunh, M., Osborne Nishimura, E., Sondek, J.E., and Dangl, J.L. (2017). TIR-only protein RBA1 recognizes a pathogen effector to regulate cell death in Arabidopsis. Proc Natl Acad Sci U S A 114, E2053–E2062.

Peart, J.R., Mestre, P., Lu, R., Malcuit, I., and Baulcombe, D.C. (2005). NRG1, a CC-NB-LRR protein, together with N, a TIR-NB-LRR protein, mediates resistance against tobacco mosaic virus. Curr Biol 15, 968–973.

Pettersen, E.F., Goddard, T.D., Huang, C.C., Meng, E.C., Couch, G.S., Croll, T.I., Morris, J.H., and Ferrin, T.E. (2021). UCSF ChimeraX: Structure visualization for researchers, educators, and developers. Protein Science 30, 70–82.

Pruitt, R.N., Locci, F., Wanke, F., Zhang, L., Saile, S.C., Joe, A., Karelina, D., Hua, C., Frohlich, K., Wan, W.L., Hu, M., Rao, S., Stolze, S.C., Harzen, A., Gust, A.A., Harter, K., Joosten, M., Thomma, B., Zhou, J.M., Dangl, J.L., Weigel, D., Nakagami, H., Oecking, C., Kasmi, F.E., Parker, J.E., and Nurnberger, T. (2021). The EDS1-PAD4-ADR1 node mediates Arabidopsis pattern-triggered immunity. Nature 598, 495–499.

Punjani, A., Rubinstein, J.L., Fleet, D.J., and Brubaker, M.A. (2017). cryoSPARC: algorithms for rapid unsupervised cryo-EM structure determination. Nat Methods 14, 290–296.

Qi, T., Seong, K., Thomazella, D.P.T., Kim, J.R., Pham, J., Seo, E., Cho, M.J., Schultink, A., and Staskawicz, B.J. (2018). NRG1 functions downstream of EDS1 to regulate TIR-NLR-mediated plant immunity in Nicotiana benthamiana. Proc Natl Acad Sci U S A 115, E10979–E10987.

Song, W., Liu, L., Yu, D.L., Bernardy, H., Jirschitzka, J., Huang, S.J., Jia, A.L., Jemielniak, W., Acker, J., Laessle, H., Wang, J.L., Shen, Q.C., Chen, W.J., Li, P.L., Parker, J.E., Han, Z.F., Schulze-Lefert, P., and Chai, J.J. (2024). Substrate-induced condensation activates plant TIR domain proteins. Nature 627, 847–853.

Sutton, R.B., Davletov, B.A., Berghuis, A.M., Sudhof, T.C., and Sprang, S.R. (1995). Structure of the first C2 domain of synaptotagmin I: a novel Ca2+/phospholipid-binding fold. Cell 80, 929–938.

Thines, B., Katsir, L., Melotto, M., Niu, Y., Mandaokar, A., Liu, G., Nomura, K., He, S.Y., Howe, G.A., and Browse, J. (2007). JAZ repressor proteins are targets of the SCF(COI1) complex during jasmonate signalling. Nature 448, 661–665.

Thor, K., Jiang, S., Michard, E., George, J., Scherzer, S., Huang, S., Dindas, J., Derbyshire, P., Leitao, N., DeFalco, T.A., Koster, P., Hunter, K., Kimura, S., Gronnier, J., Stransfeld, L., Kadota, Y., Bucherl, C.A., Charpentier, M., Wrzaczek, M., MacLean, D., Oldroyd, G.E.D., Menke, F.L.H., Roelfsema, M.R.G., Hedrich, R., Feijo, J., and Zipfel, C. (2020). The calcium-permeable channel OSCA1.3 regulates plant stomatal immunity. Nature 585, 569–573.

Tian, H., Wu, Z., Chen, S., Ao, K., Huang, W., Yaghmaiean, H., Sun, T., Xu, F., Zhang, Y., Wang, S., Li, X., and Zhang, Y. (2021). Activation of TIR signalling boosts pattern-triggered immunity. Nature 598, 500–503.

Tian, W., Wang, C., Gao, Q., Li, L., and Luan, S. (2020). Calcium spikes, waves and oscillations in plant development and biotic interactions. Nat Plants 6, 750–759.

Tian, W., Hou, C., Ren, Z., Wang, C., Zhao, F., Dahlbeck, D., Hu, S., Zhang, L., Niu, Q., Li, L., Staskawicz, B.J., and Luan, S. (2019). A calmodulin-gated calcium channel links pathogen patterns to plant immunity. Nature 572, 131–135.

Ueguchi-Tanaka, M., Ashikari, M., Nakajima, M., Itoh, H., Katoh, E., Kobayashi, M., Chow, T.Y., Hsing, Y.I., Kitano, H., Yamaguchi, I., and Matsuoka, M. (2005). GIBBERELLIN INSENSITIVE DWARF1 encodes a soluble receptor for gibberellin. Nature 437, 693–698.

Wagner, S., Stuttmann, J., Rietz, S., Guerois, R., Brunstein, E., Bautor, J., Niefind, K., and Parker, J.E. (2013). Structural basis for signaling by exclusive EDS1 heteromeric complexes with SAG101 or PAD4 in plant innate immunity. Cell Host Microbe 14, 619–630.

Wan, L., Essuman, K., Anderson, R.G., Sasaki, Y., Monteiro, F., Chung, E.H., Osborne Nishimura, E., DiAntonio, A., Milbrandt, J., Dangl, J.L., and Nishimura, M.T. (2019). TIR domains of plant immune receptors are NAD(+)-cleaving enzymes that promote cell death. Science 365, 799–803.

Wang, C., Tang, R.J., Kou, S., Xu, X., Lu, Y., Rauscher, K., Voelker, A., and Luan, S. (2024a). Mechanisms of calcium homeostasis orchestrate plant growth and immunity. Nature 627, 382–388.

Wang, H., Song, S., Gao, S., Yu, Q., Zhang, H., Cui, X., Fan, J., Xin, X., Liu, Y., Staskawicz, B., and Qi, T. (2024b). The NLR immune receptor ADR1 and lipase-like proteins EDS1 and PAD4 mediate stomatal immunity in Nicotiana benthamiana and Arabidopsis. Plant Cell 36, 427–446.

Wang, J., Hu, M., Wang, J., Qi, J., Han, Z., Wang, G., Qi, Y., Wang, H.W., Zhou, J.M., and Chai, J. (2019a). Reconstitution and structure of a plant NLR resistosome conferring immunity. Science 364, eaav5870.

Wang, J., Wang, J., Hu, M., Wu, S., Qi, J., Wang, G., Han, Z., Qi, Y., Gao, N., Wang, H.W., Zhou, J.M., and Chai, J. (2019b). Ligand-triggered allosteric ADP release primes a plant NLR complex. Science 364.

Wittig, I., Braun, H.P., and Schägger, H. (2006). Blue native PAGE. Nature Protocols 1, 418–428.

Yao, R., Ming, Z., Yan, L., Li, S., Wang, F., Ma, S., Yu, C., Yang, M., Chen, L., Chen, L., Li, Y., Yan, C., Miao, D., Sun, Z., Yan, J., Sun, Y., Wang, L., Chu, J., Fan, S., He, W., Deng, H., Nan, F., Li, J., Rao, Z., Lou, Z., and Xie, D. (2016). DWARF14 is a non-canonical hormone receptor for strigolactone. Nature 536, 469–473.

Yuan, M., Jiang, Z., Bi, G., Nomura, K., Liu, M., Wang, Y., Cai, B., Zhou, J.M., He, S.Y., and Xin, X.F. (2021). Pattern-recognition receptors are required for NLR-mediated plant immunity. Nature 592, 105–109.

Zheng, S.Q., Palovcak, E., Armache, J.P., Verba, K.A., Cheng, Y., and Agard, D.A. (2017). MotionCor2: anisotropic correction of beam-induced motion for improved cryo-electron microscopy. Nat Methods 14, 331–332.

