## Supplemental Tables for "Switch of TIR signaling by a Ca^2+^ sensor activates ADR1 recognition of pRib-AMP-EDS1-PAD4 for stomatal immunity"

**Table S1. Cryo-EM data collection, refinement and validation statistics.**

| | AtEPA complex |
|---|---|
| Data collection and processing | |
| Magnification | 130,000 |
| Voltage (kV) | 300 |
| Electron exposure ($e^-/A^2$) | 50 |
| Defocus range (μm) | -1.4~-1.8 |
| Pixel size (A) | 0.96 |
| Symmetry imposed | C2 |
| Raw movies | 14,122 |
| Particle Number | 1,779,204 |
| Map resolution (A) | 2.28 |
| FSC threshold | 0.143 |
| Map resolution range (A) | 40~2.8 |
| Refinement | |
| Protein residues | 1561 |
| Ligand | AMP:1<br>RPW:5 |
| *B* factors ($A^2$) | |
| Protein | 45.54 |
| Ligand | 20.97 |
| Water | 30.47 |
| R.m.s. deviations | |
| Bond lengths (Å) | 0.004 |
| Bond angles (°) | 0.594 |
| Validation | |
| MolProbity score | 1.33 |
| Clashscore | 3.31 |
| Ramachandran plot | |
| Favored (%) | 96.91 |
| Allowed (%) | 2.96 |
| Disallowed (%) | 0.00 |
| PDB code | 9JBN |
| EMDB code | EMD-61320 |

**Table S2. Primers used in this study.**

| **Cloning and point-mutation primers** | | |
|---|---|---|
| Recombinant Vector | Primer F | Primer R |
| *pE1776-NbSTIR1-6HA* | caaatcgactctaggggtaccATGCAACGTTCAGCAATATCTTCC | gtatgggtaagcagctctagaATAGTTGTGTAAAAGTTGTTTTCGACG |
| *pE1776-NbSTIR1-3Flag* | gccgccccctcacctctagaATGCAACGTTCAGCAATATCTTCC | atggtccttataatcactagtATAGTTGTGTAAAAGTTGTTTTCGACG |

| | | |
|---|---|---|
| *pE1776-NbSTIR1$^{E109A}$-6HA* | CTTGCATgcgTTGTCTCTGATGATGGAATTAAAGAAG | GAGACAAcgcATGCAAGCAAAAATAAGAATCACAA |
| *pE1776-NbSTIR2-6HA* | atctatctctctcgaggtaccATGCAACGTTCAGCAATATCTTCC | gtatgggtaagcagctctagaATAGTTGTGTAAAAGTTGTTTTCGACG |
| *pE1776-NbSTIR2$^{E109A}$-6HA* | CTTGCATgcgTTGTCTCTAATGATGGAATCAAAGAAG | GAGACAAcgcATGCAAGCAAAAATAAGAATCACG |
| *pCambia1300-pSTIR1:STIR1-mVenus* | acgacggccagtgccAAGCTTCGTGCCAAATTTGTTCACAACA | TGCTGAACGTTGCATGGTACCGGCCCTTTCAAGAAGTTAAAA |
| | gtaccATGCAACGTTCAGCAATATCTTCC | ggatgcggcagcagagaattcATAGTTGTGTAAAAGTTGTTTTCGACG |
| *pET-MBP-His-STIR1-GFP* | cgcggcagccatatggctagcATGCAACGTTCAGCAATATCTTCC | cttgctcactttcatggatccATAGTTGTGTAAAAGTTGTTTTCGACG |
| *pGEX4T-GST-STIR1* | gatctggttccgcgtggatccATGCAACGTTCAGCAATATCTTCC | gacgtcgtatgggtaggatccATAGTTGTGTAAAAGTTGTTTTCGACG |
| *pET-His-BdTIR* | ctgttccaggggccccatatgGCTTCTTCTGGTCTTTCCTCCA | ggtttctttaccagactcgagGAGCCTAGACAATATCATGGTTTCAC |
| *pET-His-STIR1* | ctgttccaggggccccatatgCAACGTTCAGCAATATCTTCCTCTT | ggtttctttaccagactcgagATAGTTGTGTAAAAGTTGTTTTCGACG |
| *pET-His-STIR1$^{E109A}$* | CTTGCATgcgTTGTCTCTGATGATGGAATTAAAGAAG | GAGACAAcgcATGCAAGCAAAAATAAGAATCACAA |
| *pE1776-AtEDS1-3Flag* | gccgccccccttcacctctagaATGGCGTTTGAAGCTCTTACCG | atggtccttataatcactagtGGTATCTGTTATTTCATCCATCATATAGTC |
| *pE1776-AtPAD4-3Flag* | gccgccccccttcacctctagaATGGACGATTGTCGATTCGAGA | atggtccttataatcactagtAGTCTCCATTGCGTCACTCTCAT |
| *pE1776-AtRBA1-6HA* | caaatcgactctaggggtaccATGACGAGCGTGTCTCCTCG | gtatgggtaagcagctctagaAACGGTTTGACAGTGATGATCTTC |
| *pE1776-AtEDS1-6HA* | caaatcgactctaggggtaccATGGCGTTTGAAGCTCTTACCG | gtatgggtaagcagctctagaGGTATCTGTTATTTCATCCATCATATAGTC |

| | | |
|---|---|---|
| *pE1776-StrepII-AtEDS1* | ccgcagttcgaaaaaATGGCGTTTGAAGCTCTTACCG | cacctctgttaattcgagctcCTAGGTATCTGTTATTTCATCCATCATATAGTC |
| *pE1776-AtPAD4-mGFP* | caaatcgactctaggggtaccATGGACGATTGTCGATTCGAGA | ggatgcggcagcagagaattcAGTCTCCATTGCGTCACTCTCAT |
| *pE1776-StrepII-AtPAD4* | ccgcagttcgaaaaaAAGCTTGACGATTGTCGATTCGAGAC | cacctctgttaattcgagctcCTAAGTCTCCATTGCGTCACTCTC |
| *pE1776-AtADR1-L2-3Flag* | gccgccccccttcacctctagaATGGCAGATATAATCGGCGG | atggtccttataatcactagtATCGTCGAGCCAATCCCTGCTGA |
| *pE1776-Flag-AtADR1-L2* | gacgatgacgataagaagcttGCAGATATAATCGGCGGCGAAGTT | cacctctgttaattcgagctcCTAATCGTCGAGCCAATCCCT |
| *pE1776-ISIC1-6HA* | caaatcgactctaggggtaccATGGGAAATTGCTGCTCCG | gtatgggtaagcagctctagaTAAAGTCGGCTGTATCTTTCTGAACC |
| *pE1776-ISIC1-3Flag* | gccgccccccttcacctctagaATGGGAAATTGCTGCTCCG | atggtccttataatcactagtTAAAGTCGGCTGTATCTTTCTG |
| *pET-His-ISIC1* | ctgttccaggggccccatatgGGAAATTGCTGCTCCGACG | gacgtcgtatgggtaggatccTAAAGTCGGCTGTATCTTTCTGAACC |
| *pET-His-ISIC1*$^{D\text{-}penta}$ | ctgttccaggggccccatatgGGAAATTGCTGCTCCGACG | CCATAGGgttACTCTTGGAAAGCACATCACGAT |
| | CAAGAGTaacCCTATGGCAGTCCTTTATTCAAAA | gttaacattGTACACGCGAAACACCAAATTCT |
| | GCGTGTACaatgttaacACTCAATTTCAAAATCAAGATGTCAAG | CACCAGAAATGGattACTTTTTGAGAACAAATCCTTTGATTC |
| | GTaatCCATTTCTGGTGGTCTCAAAAGC | CCGTTGCTATTGAAGTtATAACACTCGATAGTTAATGGGCTGT |
| | aACTTCAATAGCAACGGCAAGCA | gacgtcgtatgggtaggatccTAAAGTCGGCTGTATCTTTCTGAACC |
| *pE1776-AtEDS1*$^{R493A}$*-6HA* | GACCAACCgcaTACATATATGCTCAGAGAGGCTACGA | TATGTAtgcGGTTGGTCTTCCTCTTTTCATGTAC |

| | | |
|---|---|---|
| *pE1776-AtPAD4$^{R314A}$-6HA* | CAGCAgcaCTCGAGATTCAATGGTACAAAGATCG | AATCTCGAGtgcTGCTGGCAAGACACTAGCAAGC |
| *pE1776-StrepII-AtPAD4$^{K402A}$* | GCGAATTTCTACgcAAACAGAGATATAAAGACTGGCGGG | GTTTgcGTAGAAATTCGCAATGTCGAGTGGCTC |
| *pE1776-Flag-AtADR1-L2$^{D662A}$* | TTGTGcTGATCTTCTGGAACTACCTTCGACCA | CCAGAAGATCAgCACAATGATCTATTGTAAGATCAGACAAT |
| *pE1776-Flag-AtADR1-L2$^{\Delta 10}$* | CAAATCGACTCTAGGGGTACCATGGATTATAAGGACGATGACGATAA | cacctctgttaattcgagctcctaTTCCGCAGCTTCAACACGA |
| *pE1776-Flag-AtADR1-L2$^{W808A}$* | CAAATCGACTCTAGGGGTACCATGGATTATAAGGACGATGACGATAA | cacctctgttaattcgagctcCTAATCGTCGAGCgcATCCCTgctgaaagat |
| *pE1776-Flag-AtADR1-L2$^{R806A}$* | CAAATCGACTCTAGGGGTACCATGGATTATAAGGACGATGACGATAA | cacctctgttaattcgagctcCTAATCGTCGAGCCAATCcgcGCTGAAAGATTTTTCCGCAGCTTCA |
| *TRV-STIR1/2* | gtgagtaaggttaccgaattcATTATCTTGGGCTGCAGCCA | gggacatgcccgggcctcgagTGTATTTTGCCTCCTCAAGAGCC |
| *TRV-ISIC1/2* | gtgagtaaggttaccgaattcTGGCCCAGTTTCTCATTGCT | gggacatgcccgggcctcgagCGAAATACTTCCGCTCACTGTTC |
| **RT-qPCR primers** | | |
| qRT-STIR1/2 | GACGTGTTCATAAACCACAGAGG | ACTGTCCAAAAATGGCTGCAGC |
| qRT-ISIC1/2 | CAGTTTCTCATTGCTTCAACCTAAATGGT | CATTATGAAGAGCACTCATGTATGCCAT |
